# Genome-wide features of introns are evolutionary decoupled among themselves and from genome size throughout Eukarya

**DOI:** 10.1101/283549

**Authors:** Irma Lozada-Chávez, Peter F. Stadler, Sonja J. Prohaska

## Abstract

The impact of spliceosomal introns on genome and organismal evolution remains puzzling. Here, we investigated the correlative associations among genome-wide features of introns from protein-coding genes (*e.g.*, size, density, genome-content, repeats), genome size and multicellular complexity on 461 eukaryotes. Thus, we formally distinguished simple from complex multicellular organisms (CMOs), and developed the program `GenomeContent` to systematically estimate genomic traits. We performed robust phylogenetic controlled analyses, by taking into account significant uncertainties in the tree of eukaryotes and variation in genome size estimates. We found that changes in the variation of some intron features (such as size and repeat composition) are only weakly, while other features measuring intron abundance (within and across genes) are not, scaling with changes in genome size at the broadest phylogenetic scale. Accordingly, the strength of these associations fluctuates at the lineage-specific level, and changes in the length and abundance of introns within a genome are found to be largely evolving independently throughout Eukarya. Thereby, our findings are in disagreement with previous estimations claiming a concerted evolution between genome size and introns across eukaryotes. We also observe that intron features vary homogeneously (with low repetitive composition) within fungi, plants and stramenophiles; but they vary dramatically (with higher repetitive composition) within holozoans, chlorophytes, alveolates and amoebozoans. We also found that CMOs and their closest ancestral relatives are characterized by high intron-richness, regardless their genome size. These patterns contrast the narrow distribution of exon features found across eukaryotes. Collectively, our findings unveil spliceosomal introns as a dynamically evolving non-coding DNA class and strongly argue against both, a particular intron feature as key determinant of eukaryotic gene architecture, as well as a major mechanism (adaptive or non-adaptive) behind the evolutionary dynamics of introns over a large phylogenetic scale. We hypothesize that intron-richness is a pre-condition to evolve complex multicellularity.

## INTRODUCTION

Spliceosomal introns are not only germane to eukaryote origins, they also represent an evolutionary innovation on the way in which protein-coding genes have been stored, expressed and inherited throughout Life’s history on Earth. Spliceosomal introns (hereafter “introns”) form a class of non-coding DNA (ncDNA) sequences that interrupt exons within a gene. Thus, they have to be removed from the primary transcript by the splicing machinery to form a mature messenger RNA (mRNA), and hence, a functional RNA or protein molecule. Although introns are ubiquitous sequences along eukaryotes, their genome-wide features (such as length and abundance within and across genes) differ among species, and their the origins are still under intense debate. Yet, some large scale patterns of intron evolution appear to be reaching a consensus. For instance, the high conservation of intron-positions found in orthologous genes throughout eukaryotes suggests the existence of intron-rich ancestors [1, 2]. Also, intron loss has been found to be more frequent than intron gain in most lineages [1, 3–7], although episodes of rapid and extensive intron gain are also observed across eukaryotes [8–13]. Other evolutionary and functional aspects of introns remain amongst the longest-abiding puzzles, such as the phenotypic consequences of harboring intron-rich genes and their evolutionary relationship with genomic and multicellular complexity.

Because introns are less evolutionarily constrained than coding sequences, they usually evolve at high rates as do 4-fold degenerate sites and other non-coding regions [14, 15]. Nevertheless, a variable proportion of intron sites has been found to be under selective constraints in mammals [15–17], some invertebrates [18, 19], fungi[20], algae and plants [21–23]. Also, the energetic and time costs to transcribe and splice introns can be significant enough to influence the organism’s phenotype [24–27]. This is expected, in part, because eukaryotic genomes are pervasively transcribed [28, 29] and intronic RNAs constitute a major fraction of the transcribed non-coding sequences [30]. For instance, the transcription of a large gene, as the one encoding human dystrophin (2.3 Mbs), can still take up to 10 hrs. at an *in vivo* RNA pol-II elongation rate of 3.8 kb/min [31] because 99% of its length is intronic [32]. A substantial delay of gene expression owing to transcription and splicing of long and/or numerous introns, a phenomenon termed *“intron delay”* [33, 34], has turned out to be essential for cells with short mitotic cycles and for timing mechanisms during early body segmentation [35, 36]. Also, levels of gene expression (either housekeeping or tissue specific) are often associated to particular intron features [37]. For instance, some highly expressed genes are found under strong selection to remain intron-poor for transcriptional efficiency [25, 38, 39], whereas other genes are found to have longer and numerous introns to enhance expression [40–44].

Although co-transcriptional splicing depends on many parameters [45, 46], the exon-intron structure of genes is also found to have an impact on the mode of splice-site recognition and the efficiency of splicing [45, 47–51]. For instance, it was recently found that intron-containing genes and intron-rich genomes are best protected against R-loop accumulation, and subsequent transcription-associated genetic instability, by favoring spliceosome recruitment [52]. Studies across phylogenetically distant eukaryotes have also found that the length of introns and exons exerts an important influence on the likelihood of an exon to be constitutively or alternatively spliced [47–51, 53–55]. For instance, canonical splicing errors produced by splice-site recognition across small introns are more likely to result in *“Intron Retention”* or unspliced mRNA [54]. Whereas *“Exon Skipping”* or inclusion of alternative exons in the mRNA is thought to comply best with splice-site recognition across small exons [54]. Remarkably, these canonical and other noncanonical splicing errors can account for the major portion of the alternative splicing (AS) events in some eukaryotes and cell types [53, 56–58]. The expansion of AS events is thought to be key in the emergence of multicellular complexity, by creating proteome diversity and by regulating gene expression post-transcriptionally through RNA surveillance pathways [2, 59]. Accordingly, species with more tissues and cell types tend to have more alternatively spliced genes [53, 60, 61].

It remains to be fully understood, however, to what extent intron-richness (and potentially increased AS events) is coupled to the evolution of complex multicellularity (as defined in Appendix 1), genome size and of other ncDNA classes across eukaryotes. In earlier studies, a number of strong positive correlations over large evolutionary scales was found among genome size and particular ncDNA classes [62–64], including the average size, total number and nucleotide content of introns in the genome [62, 64–66]. These results have fueled the suggestion that changes in the genomic features of any ncDNA class are scaling uniformly with changes in genome size, leading to the premise that larger genomes tend to harbor more and longer introns, and *vice versa* for smaller genomes. However, correlative associations over large evolutionary scales are prone to result in biases due to low phylogenetic diversity and the lack of both phylogeny-controlled statistics [67, 68] and systematically obtained datasets. Challenging these results are also the studies showing that the average number of introns per gene (*i.e., intron density*, see Appendix 2) appears to inversely correlate with generation time [26, 69] and gene expression levels (as reviewed in [34, 37]). Accordingly, genome-wide changes of intron density are found to vary widely across eukaryotic lineages [2, 7, 70], with no clear association to genome size or another intron feature [6, 7]. Furthermore, it has been largely presumed that repeats –in particular, transposable elements (TEs)– are strongly driving the evolution of some intron features [66, 71, 72]. However, the strong contribution of repeats to intron size, for instance, is supported by studies on a few model species, particular clades or repeat families [73–76].

Some of the findings and correlative associations described previously have been interpreted as evidence for either adaptive or non-adaptive forces being the major determinant of the intron-richness complexity observed across eukaryotes. And to argue, consequently, on whether the functionality of introns can be mainly explained by the effect for which it was selected for (*i.e., selected-effects*) or by the effect of causal-role activities [29, 71, 72, 77]. For instance, the “genomic design” model postulates that the length and number of introns is determined by selection for gene function and the necessity to preserve conserved intronic elements for complex regulation [78]. Likewise, the “selection for economy” model proposes that decreases in genic size are the results of selected mutations (mainly on the length and number of introns) to reduce the time and energetic costs of transcription [25, 38, 39]. By contrast, the “mutational bias” model states that the abundance and length of introns in certain chromosomal regions is driven by different recombination rates and/or transcription-associated mutational biases [38, 70, 79]. Alternatively, the “mutation-hazard” model suggests that variation in the hazardous accumulation of introns –along with other ncDNA sequences– is primarily the outcome of increased genetic drift when effective population sizes (*N_e_*) remain small for an extended period of time [62, 63, 80]. So that, for instance, larger genomes would have more and larger introns owing to insufficient purifying selection to remove them in species with lower values of *N_e_*, a condition expected to occur in multicellular organisms particularly. The strong correlations reported among genome size, *N_e_µ* (as a proxy of *N_e_*) and some ncDNA classes –including some genome-wide features of introns– appear to support this hypothesis [62, 63, 65, 66, 68, 81]. Several controversies have emerged, however, from the contradicting evidence and arguments supporting all previous hypotheses, as discussed in [37, 67].

The discrepancy between the current observations and the evolutionary models have raised a conundrum: are the genome wide features of introns within protein-coding genes (such as their length, abundance and repetitive composition) evolving throughout Eukarya in either a concerted or an independent way among themselves, with genome size and multicellular complexity? Our study contributes to clarify this conundrum by investigating the correlative associations among these organism traits over 461 eukaryotes. To that end, we formally distinguish simple from complex multicellular organisms (CMOs) (see Appendix 1), and developed the program `GenomeContent` to systematically estimate genomic traits (see Appendix 2). We then estimated correlations under phylogenetic controlled analyses, taking into account significant uncertainties in the tree of eukaryotes and variation in genome size estimates. We found that intron features are weakly correlated among themselves and with genome size at the broadest phylogenetic scale, revealing different associations between those features estimating intron abundance across genes and those measuring intron length and repeat composition. We also found that CMOs and their closest ancestral relatives are characterized by high intron-richness, regardless their genome size. These patterns contrast the narrow distribution of exon features found across eukaryotes. Our findings are thus in disagreement with previous estimations claiming a concerted evolution between genome size and introns over a large phylogenetic scale. They also argue strongly against both a particular intron feature as key determinant of eukaryotic gene architecture, as well as a major mechanism (adaptive or non-adaptive) behind the evolutionary dynamics of introns over a large phylogenetic scale. Here, spliceosomal introns are unveiled as a dynamically evolving ncDNA class, whose relationships with genome and organismal complexity are better explained by the influence of numerous life-history factors and evolutionary forces. We argue why intron-rich lineages are more likely to evolve complex multicellularity.

## RESULTS

### Phylogenetic signal is strong in genome-based traits and robust to uncertainties in the tree of eukaryotes

*Phylogenetic signal* is the “tendency of related species to resemble each other more than species drawn at random from the same tree” [82, 83]. Because the genome traits of the 461 eukaryotes analyzed here share an evolutionary history, we first evaluated the strength of their phylogenetic signal (*i.e.*, statistical dependence) with four alternative “species trees”, each of which has been extensively used in the literature. Figure 1 shows the comparison of the tree topologies: a tree based on literature consensus from sequence-based phylogenies (Figure 1a); two NCBI taxonomy-based trees, one with no polytomies (Figure 1b), while another one with polytomies (Figure 1c); and a protein domain-based tree corrected for protein content biases derived from differences in genome size and lifestyles (Figure 1d). The literature consensus-based tree was selected as the “reference tree for eukaryotes” (Figure 1a) to present the results throughout the article. However, we do not attempt to single out any particular tree topology as the best or the correct species tree of eukaryotes. Instead, the goal is to test the robustness of the comparative analyses to alternative phylogenetic assumptions [67, 82].

**Figure 1.**
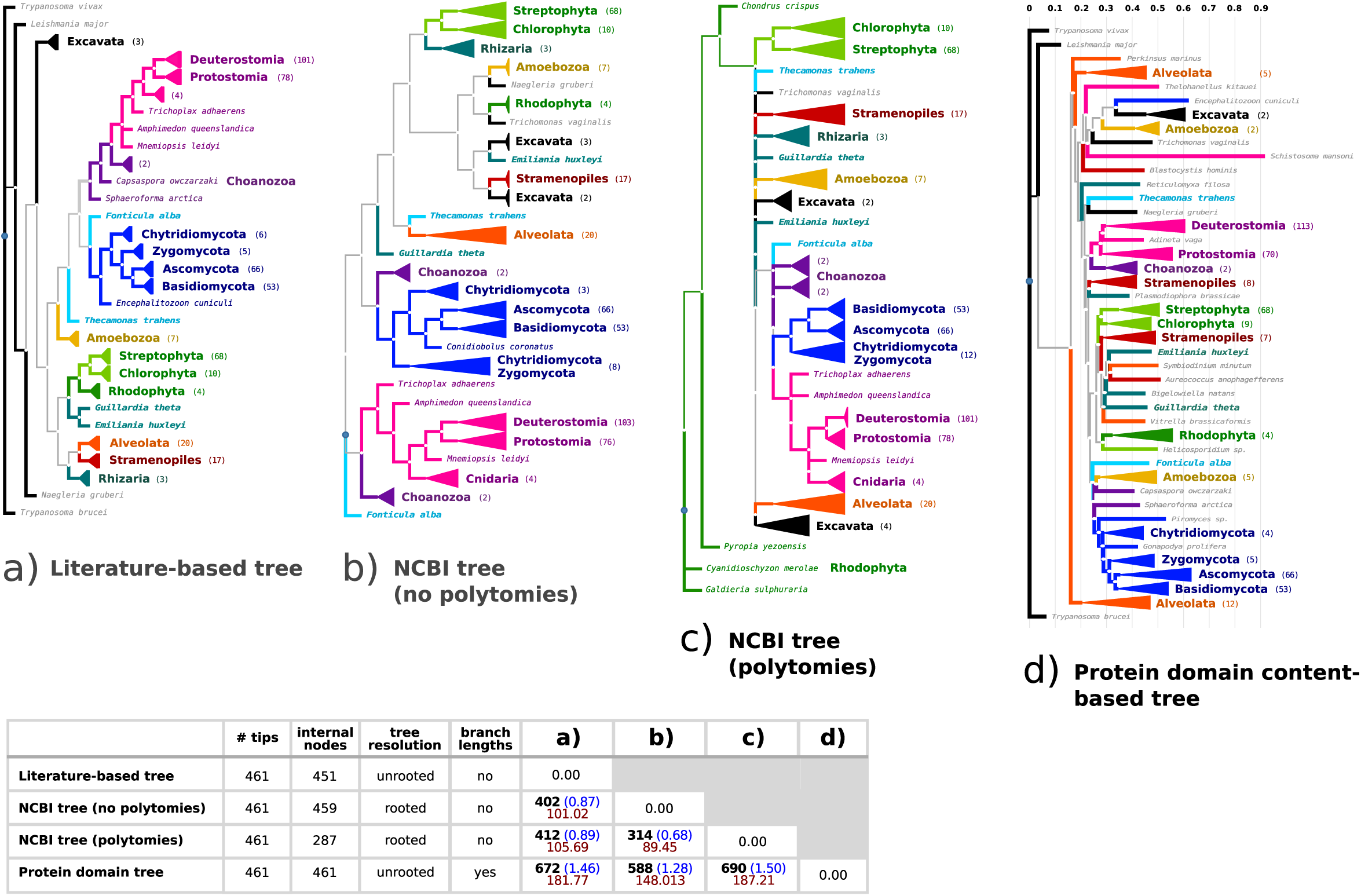
Comparison of the four alternative tree topologies estimated for the 461 eukaryotes analyzed in this study. (a) A consensus literature-based tree was manually created according to sequence-based phylogenies and supertrees. Two NCBI taxonomy-based trees were obtained in two versions: (b) not allowing polytomies and (c) allowing polytomies. (d) A protein domain content-based tree was created and corrected for protein content biases owing to differences in genome size and lifestyles. All tree topologies include: 131 fungal species (in dark blue), 78 species from Viridiplantae (in light green), 186 from Metazoa (in pink), 20 from Alveolata (in orange), 17 from Stramenopiles (in red), 7 from Excavata (in black), 7 from Amoebozoa (in yellow), 4 from Rhodophyta (in dark green), 4 from Choanozoa (in purple), 3 from Rhizaria, Cryptophyta and Haptophyta (in cyan), Fonticulidae and Apusozoa (in light blue). Phylogenetic uncertainties and disagreements among the trees are also summarized in the table (down). Numbers highlighted in bold correspond to the absolute symmetric differences (RF), the number of RF partitions per species is coloured in blue, and the Align mismatches scored in the best alignment of the branches are coloured in red.

Accordingly, dissimilarity metrics for the four tree topologies can be observed in Figure 1. The symmetric difference (RF) and the tree aligment (Align) metrics measure the number of clades not shared between two trees from either the total number of their partitions or the best alignment of their branches, respectively. Notably, the highest phylogenetic inconsistencies are observed in the protein domain-based tree: between 1.28 and 1.48 partitions per species when compared to the other trees (see table on Figure 1). This is because the protein domain-based phylogeny exhibits a *long branch attraction* (LBA) problem of several species that had undergone massive protein-domain loss, in spite of implementing a correcting factor for protein-domain content. Examples of LBA on Figure 1d include the myxosporean *Thelohanellus kitauei*, the trematode *Schistosoma mansoni*, the bdelloid rotifer *Adineta vaga* and the green algae *Helicosporidium sp*. In contrast, we observe a similar magnitude of branch dissimilarities between the literature-based tree and the NCBI-based trees, even with different polytomy resolutions: between 0.66 and 0.86 partitions per species. On closer inspection, however, the phylogenetic resolution at the level of species is not only different across the four trees, but also known conflicting hypotheses are observed for the phylogenetic positions of Rhodophyta, Rhizaria, Excavata, Stramenopiles, Alveolata, among others [84, 85]. As summarized above, the four alternative eukaryotic trees exhibit significant phylogenetic inconsistencies from one another. This allow us to incorporate adequate phylogenetic uncertainty into our comparative analyses to evaluate their sensitivity.

Table 1 presents estimates of strong phylogenetic signal for all 30 sequence-based genome traits analyzed here, as indicated by their Pagel’s *λ* values close to 1.0 and significantly > 0. Notably, the *λ* values are significantly robust to the phylogenetic disagreements shown by the alternative tree topologies. Likewise, *λ* values are significantly robust to different estimations of genome sizes and genome contents based on two sources: genome assemblies and experimental estimations (see Supplementary Table S4). The latter robustness is expected because, as also reported by Elliott and Gregory [64], the correlation between the assembled and estimated genome sizes is strong at the broadest phylogenetic scale: *r* = 0.958 (see Table 3). Consistent with previous studies [67, 76, 81, 86], these results indicate that genome traits are not statisticallly independent when compared among species. Therefore, correction for phylogenetic signal is accounted for any comparative analyses in this study.

### Genome size correlates weakly with genome-wide intron features at the broadest phylogenetic scale

Table 2 shows the estimated log Bayes Factors (log BF) and coefficients of determination (*r*^2^) used here as the criterion to assess both the “strength of the evidence” and the “explanatory power” of the correlative associations between two traits (*X* and *Y*), respectively. The “explanatory power” of the *r*^2^ values should be understood as means of a statistical range to associate the variation observed between *X* and *Y*, with no implication as to the evolutionary mechanism that might cause (or not) such associations nor the primary trait (*X* or *Y*) subject to this action.

**Table 1.** Unvariate measures of phylogenetic signal (with the *λ* parameter) for the genome traits analysed in this study, by using four alternate trees for the 461 eukaryotic species (see Figure 1).

| Tree topology |  | Reference |  |  | Protein | NCBI | NCBI | D.F. |
| --- | --- | --- | --- | --- | --- | --- | --- | --- |
| Genome trait | $\lambda$ | ln ML | Estimated AIC | CI (95%) | domain $\lambda$ | No Poly $\lambda$ | Yes Poly $\lambda$ | |
| Estimated genome size (Mbs) | **0.938** | -129.197 | 260.395 | (0.904, 0.960) | 0.999 | 0.945 | 0.939 | 460 |
| Assembled genome size (Mbs) | **0.936** | -117.203 | 236.406 | (0.901, 0.958) | 0.999 | 0.943 | 0.937 | 460 |
| Genome repeat content (Mbs) | **0.898** | -362.153 | 726.305 | (0.843, 0.936) | 0.999 | 0.916 | 0.889 | 460 |
| Genome repeat content (%) | **0.773** | -116.277 | 234.554 | (0.661, 0.854) | 0.965 | 0.786 | 0.757 | 460 |
| Unique genome content (Mbs) | **0.939** | -41.127 | 84.255 | (0.907, 0.960) | 0.999 | 0.938 | 0.937 | 460 |
| Unique genome content (%) | **0.759** | 421.403 | -840.806 | (0.632, 0.851) | 0.865 | 0.827 | 0.748 | 460 |
| Unique nc-genome content (Mbs) | **0.948** | -140.983 | 283.965 | (0.918, 0.967) | 0.999 | 0.929 | 0.944 | 460 |
| Unique nc-genome content (%) | **0.805** | 342.896 | -683.792 | (0.677, 0.890) | 0.971 | 0.679 | 0.755 | 460 |
| CDS number | **0.914** | 154.015 | -306.030 | (0.863, 0.947) | 0.999 | 0.896 | 0.929 | 460 |
| CDS avg size (nts) | **0.873** | 118.854 | -235.708 | (0.811, 0.917) | 0.999 | 0.924 | 0.891 | 460 |
| CDS genome coverage (Mbs) | **0.856** | -55.893 | 113.786 | (0.790, 0.904) | 0.999 | 0.847 | 0.840 | 460 |
| Exon density | **0.953** | 265.697 | -529.395 | (0.917, 0.973) | 0.999 | 0.937 | 0.940 | 460 |
| Exon number | **0.954** | 12.167 | -22.335 | (0.926, 0.972) | 0.999 | 0.924 | 0.951 | 460 |
| Exon avg-size (nts) | **0.987** | 316.981 | -631.963 | (0.979, 0.992) | 0.999 | 0.979 | 0.990 | 460 |
| Exon content (Mbs) | **0.859** | 168.034 | -334.068 | (0.776, 0.914) | 0.999 | 0.835 | 0.885 | 460 |
| Exon content (%) | **0.913** | -67.455 | 136.910 | (0.865, 0.945) | 0.999 | 0.924 | 0.887 | 460 |
| Unique exon content (Mbs) | **0.880** | 210.273 | -418.547 | (0.807, 0.926) | 0.999 | 0.851 | 0.883 | 460 |
| Unique exon content (%) | **0.479** | 742.122 | -1482.244 | (0.312, 0.637) | 0.743 | 0.500 | 0.634 | 460 |
| Repeat exon content (Mbs) | **0.797** | -244.207 | 490.415 | (0.688, 0.874) | 0.984 | 0.827 | 0.842 | 460 |
| Repeat exon content (%) | **0.776** | -141.757 | 285.515 | (0.655, 0.863) | 0.955 | 0.818 | 0.798 | 460 |
| Number of CDS with introns | **0.955** | -128.310 | 258.620 | (0.935, 0.969) | 0.999 | 0.919 | 0.950 | 460 |
| CDS with introns (%) | **0.949** | -0.495 | 2.989 | (0.929, 0.964) | 0.861 | 0.892 | 0.928 | 460 |
| Intron number | **0.960** | -222.858 | 447.715 | (0.940, 0.973) | 0.999 | 0.930 | 0.958 | 460 |
| Intron density | **0.934** | 286.690 | -571.380 | (0.890, 0.962) | 0.999 | 0.943 | 0.926 | 460 |
| Intron wm-size (nts) | **0.875** | -10.518 | 23.035 | (0.812, 0.919) | 0.999 | 0.922 | 0.875 | 460 |
| Intron content (Mbs) | **0.950** | -310.428 | 622.856 | (0.924, 0.967) | 0.999 | 0.901 | 0.948 | 460 |
| Intron content (%) | **0.932** | -174.617 | 351.234 | (0.895, 0.957) | 0.999 | 0.871 | 0.925 | 460 |
| Unique intron content (Mbs) | **0.955** | -290.605 | 583.211 | (0.931, 0.970) | 0.999 | 0.901 | 0.951 | 460 |
| Unique intron content (%) | **0.616** | 527.481 | -1052.962 | (0.447, 0.748) | 0.998 | 0.642 | 0.611 | 460 |
| Repeat intron content (Mbs) | **0.885** | -452.997 | 907.994 | (0.829, 0.925) | 0.999 | 0.926 | 0.873 | 456 |
| Repeat intron content (%) | **0.728** | -158.590 | 319.180 | (0.596, 0.824) | 0.975 | 0.666 | 0.657 | 456 |

**Table 2.**
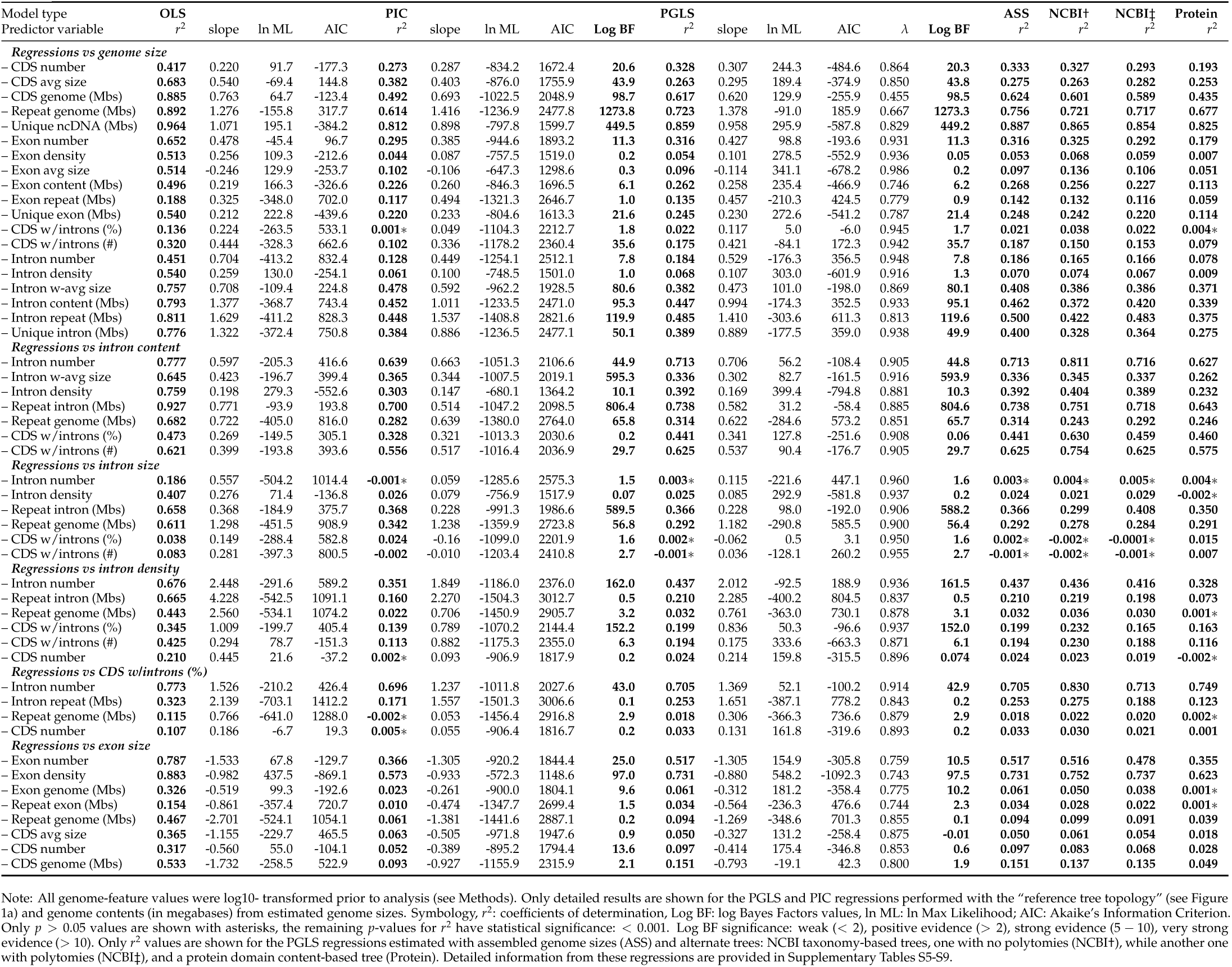
Correlative associations among several features measuring genome complexity and intron-richness from 461 eukaryotes under non-phylogenetic (OLS) and phylogenetic (PIC, PGLS) models.

**Table 3.** Summary statistics and PGLS regressions performed between genome size and intron features for different species datasets.

| Genome features<br>Taxon | *n* | GENOME:<br>size (EST vs ASS)<br>*r*<sup>2</sup> [Mbs] | repeats<br>[%] | INTRONS:<br>% in CDS<br>*r*<sup>2</sup> [%] | density<br>*r*<sup>2</sup> [no.] | size<br>*r*<sup>2</sup> [nts] | content<br>*r*<sup>2</sup> [%] | repeats<br>*r*<sup>2</sup> [%] | number<br>*r*<sup>2</sup> [no.] |
| --- | --- | --- | --- | --- | --- | --- | --- | --- | --- |
| **Eukarya** | 461 | **0.958** [—] | — | **0.022** [—] | **0.068** [—] | **0.382** [—] | **0.447** [—] | **0.485** [—] | **0.184** [—] |
| *Random and selected datasets* |  |  |  |  |  |  |  |  |  |
| **Random set 1**⊕ | 231 | **0.986** [—] | — | **0.022** [—] | **0.086** [—] | **0.343** [—] | **0.424** [—] | **0.564** [—] | **0.184** [—] |
| **Random set 2**⊕ | 116 | **0.987** [—] | — | **0.048** [—] | **0.162** [—] | **0.209** [—] | **0.526** [—] | **0.382** [—] | **0.259** [—] |
| **Random set 3**⊕ | 58 | **0.995** [—] | — | **0.197** [—] | **0.505** [—] | **0.687** [—] | **0.899** [—] | **0.723** [—] | **0.470** [—] |
| **Random set 4**⊕ | 29 | **0.992** [—] | — | **0.172** [—] | **0.481** [—] | **0.808** [—] | **0.904** [—] | **0.728** [—] | **0.291** [—] |
| **Lynch & Conery 2003**† | 26 | — | — | **0.189** [—] | **0.102** * [—] | **0.683** [—] | **0.549** [—] | **0.489** [—] | **0.300** [—] |
| **Wu & Hurst 2015**‡ | 30 | — | — | **0.214** [—] | **0.713** [—] | **0.850** [—] | **0.932** [—] | **0.895** [—] | **0.412** [—] |
| *Local phylogenetic scale* |  |  |  |  |  |  |  |  |  |
| **Protists** | 66 | **0.961** [83.6] | 21.2 | **0.024** * [56.6] | **0.181** [3.5] | **0.271** [227.3] | **0.490** [9.6] {10.3} | **0.475** [18.2] | **0.267** [42,957] |
| Stramenopiles | 17 | **0.982** [86.0] | 26.7 | **-0.062** * [59.3] | **-0.040** * [2.9] | **0.003** * [227.8] | **0.397** [7.5] {7.7} | **0.580** [19.8] | **0.330** [29,148] |
| Alveolata | 20 | **0.977** [116.1] | 12.1 | **0.248** [63.4] | **0.697** [4.5] | **0.344** [193.4] | **0.803** [11.1] {13.0} | **0.704** [14.6] | **0.481** [68,920] |
| **Fungi** | 131 | **0.985** [38.2] | 13.6 | **0.080** [65.9] | **0.034** [3.2] | **0.089** [120.4] | **0.285** [7.0] {7.0} | **0.528** [8.8] | **0.220** [32,768] |
| Basidiomycota | 53 | **0.993** [47.3] | 17.0 | **-0.001** * [81.8] | **-0.020** * [4.7] | **0.426** [92.2] | **0.598** [10.4] {10.3} | **0.694** [11.2] | **0.569** [57,516] |
| Ascomycota: | 66 | **0.964** [30.8] | 10.2 | **0.489** [53.4] | **0.218** [1.9] | **-0.009** * [141.4] | **0.641** [4.0] {4.0} | **0.633** * [6.2] | **0.193** [13,311] |
| – Pezizomycotina | 42 | **0.982** [40.6] | 13.4 | **0.018** * [72.8] | **0.147** [2.3] | **0.044** * [107.2] | **0.107** [5.2] {5.2} | **0.182** [7.2] | **-0.012** * [19,740] |
| – Saccharomycotina | 22 | **0.659** [13.7] | 4.6 | **0.092** * [16.3] | **0.145** [1.3] | **0.001** * [211.7] | **0.248** [1.7] {1.7} | **0.361** [4.5] | **0.133** * [1,661] |
| **Chlorophyta** | 10 | **0.953** [49.3] | 11.2 | **0.621** [64.7] | **0.807** [4.7] | **0.629** [245.8] | **0.939** [19.3] {19.9} | **0.806** [15.2] | **0.879** [46,269] |
| **Streptophyta** | 68 | **0.967** [1,065.8] | 41.6 | **-0.015** * [74.9] | **0.137** [4.9] | **0.415** [562.6] | **0.435** [11.8] {13.6} | **0.299** [20.2] | **0.104** [134,777] |
| Monocots | 15 | **0.978** [1,809.8] | 49.0 | **-0.024** * [75.3] | **-0.044** * [4.8] | **-0.040** * [810.9] | **-0.035** * [9.4] {11.4} | **-0.077** * [19.5] | **-0.028** * [129,949] |
| Eudicots | 49 | **0.952** [641.1] | 43.8 | **-0.018** * [76.5] | **-0.016** * [5.0] | **0.442** [495.6] | **0.537** [11.3] {13.5} | **0.484** [20.4] | **0.214** [155,964] |
| **Metazoa** | 186 | **0.973** [1,260.9] | 26.5 | **0.048** [87.0] | **-0.001** * [7.5] | **0.683** [2,643.5] | **0.678** [24.6] {26.4} | **0.690** [24.2] | **0.077** [120,238] |
| Protostomia: | 78 | **0.957** [517.9] | 25.7 | **0.138** [84.4] | **-0.011** * [5.2] | **0.659** [1,535.0] | **0.606** [21.7] {22.8} | **0.693** [24.0] | **0.104** [75,745] |
| – Lophotrochozoa | 10 | **0.914** [765.6] | 35.8 | **0.119** * [79.1] | **0.293** * [6.2] | **0.319** * [1,671.4] | **0.577** [21.5] {23.6} | **0.502** [33.1] | **-0.125** * [126,857] |
| – Arthropoda: | 61 | **0.948** [523.3] | 25.4 | **0.053** [84.3] | **-0.013** * [4.9] | **0.684** [1,640.0] | **0.595** [21.0] {22.1} | **0.657** [23.8] | **0.080** [64,879] |
| – Diptera | 13 | **0.938** [410.1] | 30.6 | **-0.090** * [84.0] | **0.186** * [3.5] | **0.738** [1,484.9] | **0.482** [14.6] {16.9} | **0.754** [28.8] | **-0.090** * [40,508] |
| – Hymenoptera | 18 | **0.646** [278.6] | 18.0 | **0.154** * [85.4] | **-0.058** * [5.3] | **-0.017** * [1,259.4] | **-0.043** * [25.4] {27.8} | **-0.009** * [15.2] | **-0.039** [59,118] |
| – Lepidoptera | 13 | **0.975** [402.0] | 29.2 | **-0.089** * [86.5] | **0.032** * [5.6] | **0.346** [1,377.7] | **0.044** * [23.9] {23.9} | **0.412** [30.4] | **-0.038** * [80,139] |
| Deuterostomia: | 101 | **0.955** [1,898.6] | 26.6 | **0.083** [90.1] | **0.077** [9.3] | **0.729** [3,623.5] | **0.728** [26.8] {29.4} | **0.614** [23.9] | **0.044** [156,583] |
| – Teleostei | 17 | **0.928** [871.6] | 20.8 | **0.324** [94.2] | **-0.009** * [9.7] | **0.856** [1,662.4] | **0.844** [31.5] {35.1} | **0.922** [19.2] | **0.348** [200,011] |
| – Amniota: | 72 | **0.567** [2,234.2] | 26.4 | **0.341** [89.6] | **0.099** [9.5] | **0.184** [4,321.7] | **0.667** [26.3] {28.8} | **0.476** [23.0] | **0.112** [149,821] |
| – Reptiles | 11 | **0.806** [2,188.3] | 33.9 | **-0.068** * [89.8] | **0.036** * [8.8] | **0.363** [4,982.6] | **0.584** [26.5] {28.2} | **0.329** [30.6] | **-0.052** * [143,335] |
| – Aves | 28 | **-0.029** * [1,231.1] | 8.5 | **0.032** * [92.9] | **-0.036** * [10.1] | **-0.028** * [3,334.5] | **-0.023** * [31.5] {33.6} | **-0.028** * [7.0] | **0.032** * [139,985] |
| – Mammalia | 33 | **0.482** [3,100.6] | 39.0 | **-0.032** * [86.8] | **-0.020** * [9.3] | **0.092** [4,938.9] | **0.169** [21.9] {24.9} | **0.377** [34.1] | **-0.007** * [160,330] |

Note: ⊕ Randomly reduced datasets from the original 461 species analyzed in this study, with 100 replicates each (see Methods). † Complete genome sequences are only available for 26 from the 32 metazoan species included in the original dataset (see Supplementary Table S10). ‡ This dataset has an overrepresentation of metazoan species (around two thirds), particularly of vertebrates (see Supplementary Table S10). All genome-feature values were log10- transformed prior to analysis (see Methods). The [mean value] for selected genome features is shown, and for genome contents (in % or Mbs) as calculated from estimated genome sizes. The *r*<sup>2</sup> values (highlighted in bold font) obtained from PGLS regressions were performed with estimated genome sizes and the “reference tree topology” (in Figure 1a). Unique or non-repetitive intron contents are denoted with {}. Only *p* > 0.05 values are shown with asterisks, the remaining *p*-values for *r*<sup>2</sup> have statistical significance: < 0.001. The regressions performed with alternative tree topologies and assembled genome sizes are provided in Supplementary Table S12. Summary statistics for additional clades are provided in Supplementary Tables S14-S15.

When the data is analyzed as phylogenetically independent with the OLS model, strong associations are observed on Table 2 among genome size and most intron features (*r*^2^ between 0.6 and 0.9). In particular, the regression between genome and intron sizes (*r*^2^ = 0.76) is consistent with previous estimations performed over a broad evolutionary range by Vinogradov (*r*^2^ = 0.792, *n* = 27) [65] and by Lynch and Conery (*r*^2^ = 0.641, *n* = 30) [62]. However, the strength of such associations substantially dropped after phylogenetic corrected regressions were performed with both PGLS and PICs models (see Table 2). For instance, the correlations among genome size and the features estimating intron length and genomic content have robust evidence for a positive association but a weak explanatory power at the broadest phylogenetic scale: *r*^2^ = 0.382 (*logBF* = 80.1) for intron size, *r*^2^ = 0.447 (*logBF* = 95.1) for intron content, and *r*^2^ = 0.485 (*logBF* = 119.6) for the repetitive-intronic content of the genome. On the other hand, none or no simple associations were found among genome size and those features measuring the abundance of introns within and across genes: *r*^2^ = 0.068 (*logBF* = 1.3) for intron density, *r*^2^ = 0.022 (*logBF* = 1.7) for the fraction (and total number) of intron-containing genes per genome, and *r*^2^ = 0.184 (*logBF* = 7.8) for the total number of introns per genome. Therefore, only reduced and differential fractions of the variation observed among intron features (from 2% to 45%) can be associated (directly or indirectly) to the ∼2,200-fold variation of the genome size observed over the 461 eukaryotes analyzed here.

The strenght of these correlations, either through log BF or *r*^2^ values, is not only similar regardless which genome trait (*e.g.*, genome size) is associate to the *X* and *Y* variables, but it is also consistent with both PGLS and PIC models (see Supplementary Tables S5-S10). Nonetheless, the PGLS model appears to fit the data significantly better than the OLS and PIC models do, as shown by its consistent lower AIC values and higher ML values (see Table 2). We further investigated potential discrepancies in our correlations owing to the differences to estimate tree topologies, genome traits, species number and diversity (see Supplementary Tables S5-S10). As shown in Table 2, the weak associations found among genome size and intron features also proved to be robust to the phylogenetic inconsistencies of the alternate trees. A slighter strenght should be notice in the correlations performed with the protein-domain content tree, which exhibits the highest topological and branch length dissimilarities in comparison to the other trees. We also found no major discrepancies in the PGLS correlations performed with estimates of intron features based on (and against) the two sources of genome size. The PGLS correlations described in Table 3 further show that the weak associations found at the broadest phylogenetic scale, significant *r*^2^ values from 0.02 to 0.5, are robust to randomly reduced datasets of 231 and 116 species sampled from the original 461 species (see also Supplementary Table S11). These results support the robustness of the phylogenetic diversity of our species dataset. By contrast, we found overestimated correlations among intron features and genome size when smaller and less diverse datasets are used to cover large phylogenetic scales. This is observed in Table 3 for two further randomly reduced datasets of 58 and 29 species, as well as for the species datasets from Lynch and Conery [62] (*n* = 26) and from Wu and Hurst [68] (*n* = 30) (see also Supplementary Table S12). These contrasting results show that phylogenetic diversity and phylogenetic corrected correlations over large evolutionary scales are strongly affected by very small and biased datasets.

Under the “replicated co-distribution” approach [87], we also tested the decoupled association between intron features and genome size across multiple independent clades. Table 3 describes the PGLS correlations performed over 18 lineage-specific datasets compiled from the original dataset of 461 eukaryotes (see Methods and Supplementary Table S11). According to our previous results, the *r*^2^ values show that the strength of the associations among genome size and intron features is indeed different at the local phylogenetic scale. For instance, the genome-intron size relationship varies from *r*^2^ = 0.003 in stramenopiles up to *r*^2^ = 0.856 in teleosts, whereas the association between intron density and genome size varies from *r*^2^ = 0.077 in deuterostomes up to *r*^2^ = 0.807 in chlorophytes. Likewise, a differing correlative association among intron features and genome size is observed in Hymenoptera, Aves, Monocots and Ascomycota when compared to the correlations observed in their corresponding close relatives.

Nevertheless, some of the correlations on Table 3 also highlight the impact that differences in the estimations of genome features have at the local evolutionary scale. For instance, the correlation obtained for genome size and intron density in Ascomycota is consistent with Kelkar and Ochman [81], regardless both the tree topology or the source for the genome size estimates used to perform the regressions. This is expected because the assembled and estimated genome sizes are highly correlated in this group (*r*^2^ = 0.964, *p* < 0.001, Table 3). By contrast, our correlations for genome and intron size in amniotes are not consistent with those reported by Zhang and Edwards [76] when the estimated genome sizes are used to perform the regressions (*r*^2^ = 0.184, *p* < 0.001, Table 3), but they are when the regressions are performed with the assembled genome sizes (*r*^2^ = 0.339, *p* < 0.001, Supplementary Table S11). Such discrepancy might be the consequence of the low correlation found between the two sources of genome size estimates in amniotes: *r*^2^ = 0.567 (*p* < 0.001, see Table 3). On the other hand, some of the regressions obtained across specific lineages (such as monocots and aves) or for particular intron features (such as intron density and the fraction intron-containing genes) do not reach statistical significance. While most of these correlations reveal none or no simple associations among intron features and genome size, caution should be taken in the correlations obtained for those lineages with few sequenced genomes (*n* < 15). With few exceptions, the PGLS correlations reported previously, and in the following sections, do not pose significant changes when they are performed with different tree topologies and assembled genome sizes (see Supplementary Tables S5-S13).

### Changes of intronic content across lineages are not strongly associated to one particular genome-wide intron feature

As observed on Figure 2 and Table 3, the net nucleotide coverage of introns in the eukaryotic genome, *i.e.* “intron content”, represents on average 25.8% of the genome size in Choanozoa, 21.9% in Metazoa, 11.8% in Viridiplantae, 11.1% in Alveolata, 10.4% in Amoebozoa, 7.5% in Stramenopiles, 7.0% in Fungi, and 0.9% in Excavata. Our results show that the variation of intron content observed at the broadest phylogenetic scale is positively, yet weakly correlated with genome size (*r*^2^ = 0.447, *logBF* = 95.1), in contrasts to the stronger associations observed with the repetitive (*r*^2^ = 0.723, *logBF* = 1273.3), non-repetitive ncDNA (*r*^2^ = 0.859, *logBF* = 449.2) and protein-coding (*r*^2^ = 0.617, *logBF* = 98.5) contents of the genome (see Table 2). As a consequence, low intron-contents are observed in some large and highly repetitive genomes, such as *Pinus taeda* (1.69% in 22.5 Gbs) and *Locusta migratoria* (13.61% in 6.5 Gbs). And *vice versa*, high non-repetitive intron contents can be observed in species with smaller genome sizes, either unicellular or multicellular, when compared to their corresponding close relatives. Some remarkable examples include Chlorophytes (29.2%, 70.7 Mbs), Aves (31.5%, 1.23 Gbs), teleosts (31.5%, 871.6 Mbs), bees (25.4%, 278.6 Mbs), buterflies (23.9%, 402.0 Mbs), *Bigelowiella natans* (32.05%, 94.7 Mbs), *Ectocarpus siliculosus* (36.04%, 214 Mbs), and *Utricularia gibba* (18.15%, 88 Mbs) (see Figures 2-4 and Supplementary Tables S14-S15). Consistent with this, we further found that the strength of the association between intron content and genome size is indeed different across lineages (Table 3). For instance, it is strong in Alveolata, Chlorophyta and Teleostei, while weak or absent in Pezizomycotina, Aves, Mammalia, Monocots, Hymenoptera and Lepidoptera. In agreement with a previous study [64], these findings show that intron content cannot fully account for the large variations of eukaryotic genome sizes.

**Figure 2.**
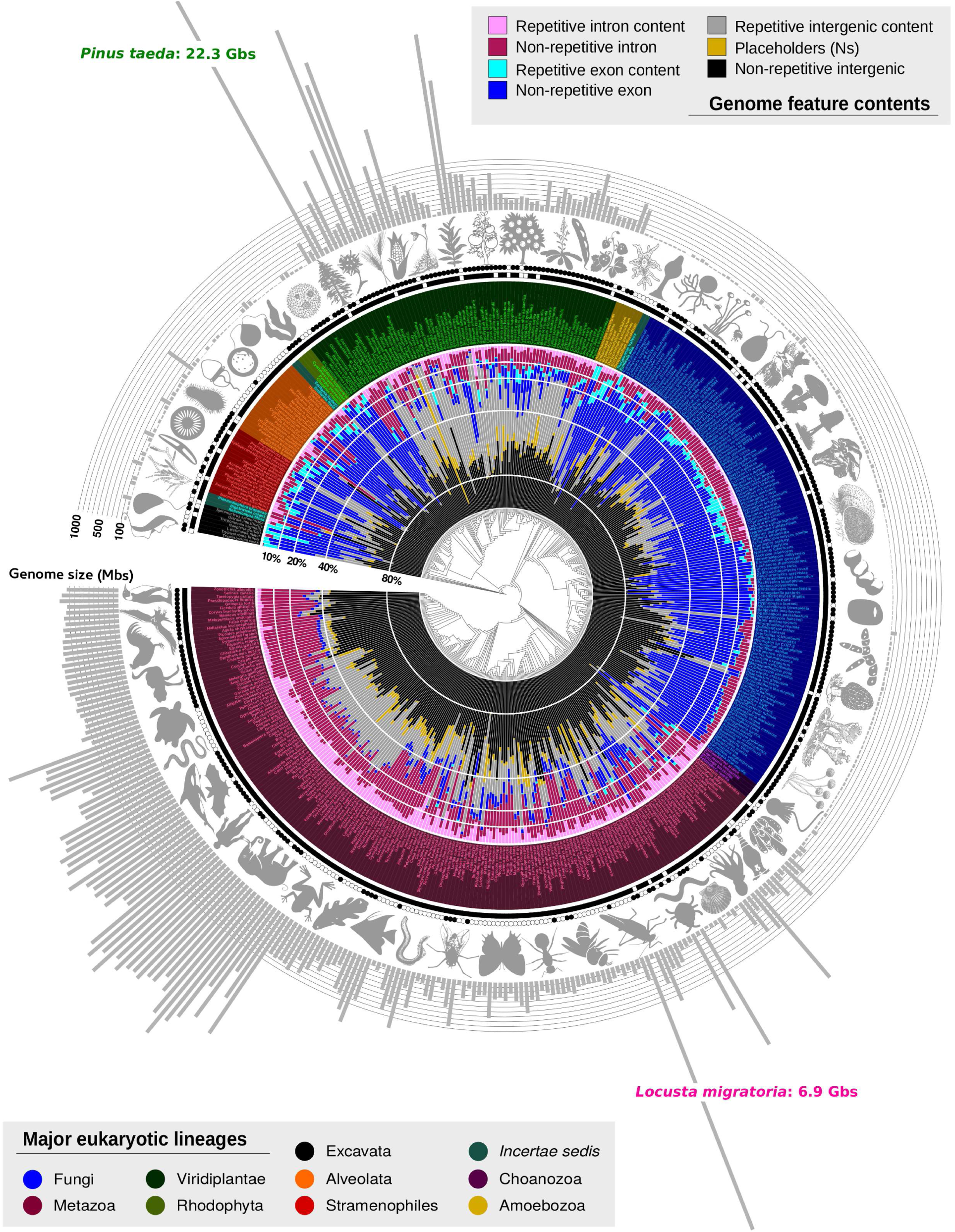
Approximate estimation of genome size and contents across 461 eukaryotes. The species are displayed according to the “reference tree for eukaryotes” (in Figure 1a) and coloured according to the supergroup they belong to. The fraction of genomic content (repetitive and non-repetitive) from the total assembled genome size –as calculated with the `GenomeContent` program–, for introns, exons and intergenic regions is coloured in red, blue and gray scales, respectively. The fraction de placeholders (sequences of Ns) is represented in yellow. Noteworthy, the fraction of non-repetitive intergenic regions might be smaller due to the presence of several repetitive pseudogenes and non-coding RNA families (*e.g.*, ribosomal RNAs, tRNAs) that are not fully annotated in the genome projects nor in the present study. The paired symbols for each species indicate whether the nucleotide overlap of repeats within intronic (circles) and exonic sequences (squares) is less (TRUE: filled) or more (FALSE: not filled) significant than expected by chance, according to the *p* < 0.05 estimated over 1,000 permutation tests on the *Jaccard index* for each feature and genome (see Methods and Supplementary Table S16). The assembled genome size for every species is shown in vertical bars. Data calculated from estimated genome sizes are available in Supplementary Table S3.

Our findings also show that intron content in eukaryotes is not strongly associated to any other particular intron feature at the broadest phylogenetic scale. As observed in Table 2, changes of intron size (*r*^2^ = 0.336), intron density (*r*^2^ = 0.392), the fraction of genes harboring introns (*r*^2^ = 0.441) and the repetitive-genome content (*r*^2^ = 0.314) are similarly associated to the variation of intron content observed across eukaryotes. To a larger extent, intron content is associated with changes in the total number of introns per genome (*r*^2^ = 0.713) and the repetitive-intron content (*r*^2^ = 0.738). It is important to clarify, however, that the strength of the association among intron content and other genome-wide intron features is different across lineages (see Table 4 and Supplementary Table S13), as we report next.

**Table 4.**
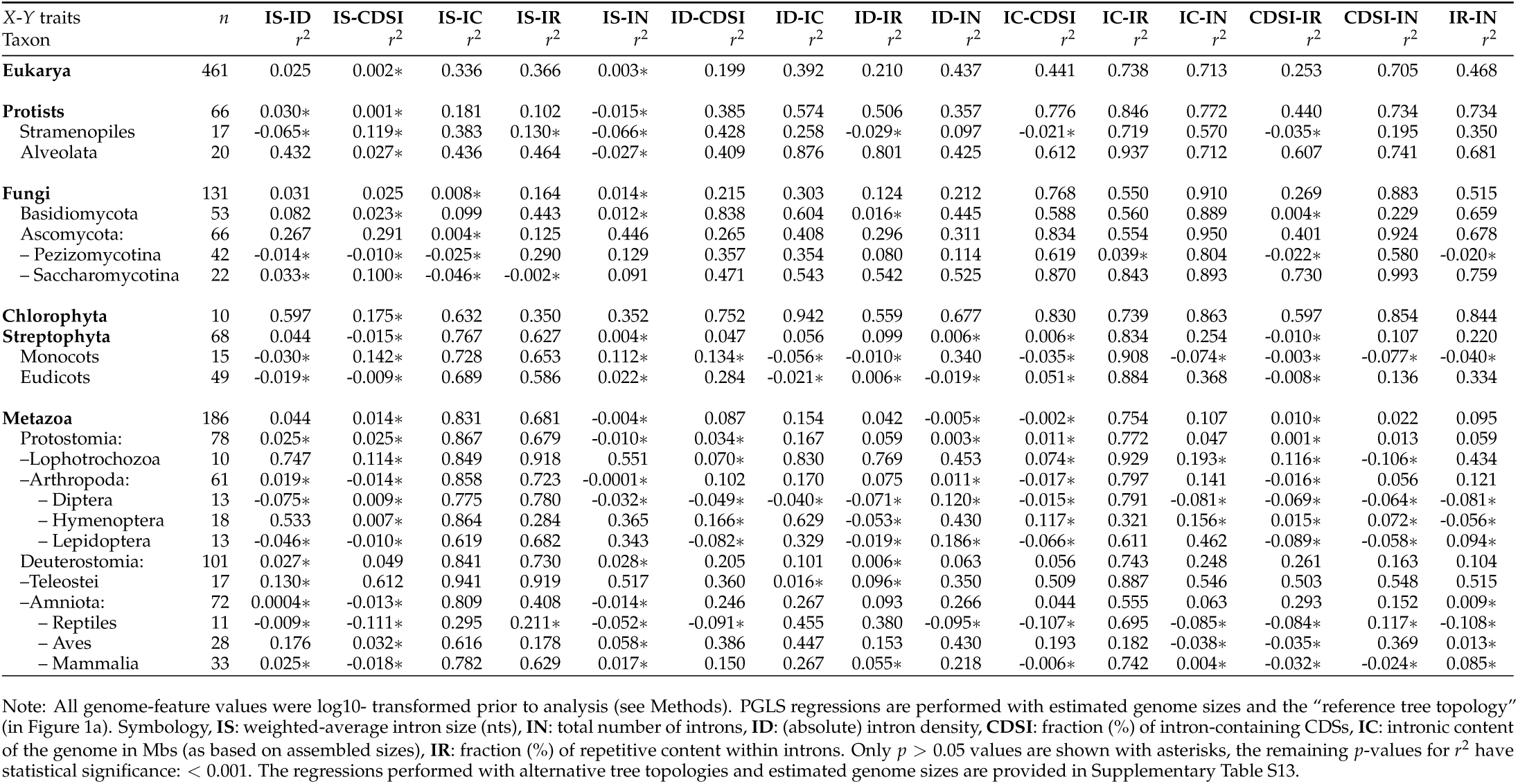
PGLS regressions performed among intron features for different taxa datasets.

| X-Y traits<br>Taxon | *n* | IS-ID<br>*r*<sup>2</sup> | IS-CDSI<br>*r*<sup>2</sup> | IS-IC<br>*r*<sup>2</sup> | IS-IR<br>*r*<sup>2</sup> | IS-IN<br>*r*<sup>2</sup> | ID-CDSI<br>*r*<sup>2</sup> | ID-IC<br>*r*<sup>2</sup> | ID-IR<br>*r*<sup>2</sup> | ID-IN<br>*r*<sup>2</sup> | IC-CDSI<br>*r*<sup>2</sup> | IC-IR<br>*r*<sup>2</sup> | IC-IN<br>*r*<sup>2</sup> | CDSI-IR<br>*r*<sup>2</sup> | CDSI-IN<br>*r*<sup>2</sup> | IR-IN<br>*r*<sup>2</sup> |
| --- | --- | --- | --- | --- | --- | --- | --- | --- | --- | --- | --- | --- | --- | --- | --- | --- |
| **Eukarya** | 461 | 0.025 | 0.002* | 0.336 | 0.366 | 0.003* | 0.199 | 0.392 | 0.210 | 0.437 | 0.441 | 0.738 | 0.713 | 0.253 | 0.705 | 0.468 |
| **Protists** | 66 | 0.030* | 0.001* | 0.181 | 0.102 | -0.015* | 0.385 | 0.574 | 0.506 | 0.357 | 0.776 | 0.846 | 0.772 | 0.440 | 0.734 | 0.734 |
| Stramenopiles | 17 | -0.065* | 0.119* | 0.383 | 0.130* | -0.066* | 0.428 | 0.258 | -0.029* | 0.097 | -0.021* | 0.719 | 0.570 | -0.035* | 0.195 | 0.350 |
| Alveolata | 20 | 0.432 | 0.027* | 0.436 | 0.464 | -0.027* | 0.409 | 0.876 | 0.801 | 0.425 | 0.612 | 0.937 | 0.712 | 0.607 | 0.741 | 0.681 |
| **Fungi** | 131 | 0.031 | 0.025 | 0.008* | 0.164 | 0.014* | 0.215 | 0.303 | 0.124 | 0.212 | 0.768 | 0.550 | 0.910 | 0.269 | 0.883 | 0.515 |
| Basidiomycota | 53 | 0.082 | 0.023* | 0.099 | 0.443 | 0.012* | 0.838 | 0.604 | 0.016* | 0.445 | 0.588 | 0.560 | 0.889 | 0.004* | 0.229 | 0.659 |
| Ascomycota: | 66 | 0.267 | 0.291 | 0.004* | 0.125 | 0.446 | 0.265 | 0.408 | 0.296 | 0.311 | 0.834 | 0.554 | 0.950 | 0.401 | 0.924 | 0.678 |
| – Pezizomycotina | 42 | -0.014* | -0.010* | -0.025* | 0.290 | 0.129 | 0.357 | 0.354 | 0.080 | 0.114 | 0.619 | 0.039* | 0.804 | -0.022* | 0.580 | -0.020* |
| – Saccharomycotina | 22 | 0.033* | 0.100* | -0.046* | -0.002* | 0.091 | 0.471 | 0.543 | 0.542 | 0.525 | 0.870 | 0.843 | 0.893 | 0.730 | 0.993 | 0.759 |
| **Chlorophyta** | 10 | 0.597 | 0.175* | 0.632 | 0.350 | 0.352 | 0.752 | 0.942 | 0.559 | 0.677 | 0.830 | 0.739 | 0.863 | 0.597 | 0.854 | 0.844 |
| **Streptophyta** | 68 | 0.044 | -0.015* | 0.767 | 0.627 | 0.004* | 0.047 | 0.056 | 0.099 | 0.006* | 0.006* | 0.834 | 0.254 | -0.010* | 0.107 | 0.220 |
| Monocots | 15 | -0.030* | 0.142* | 0.728 | 0.653 | 0.112* | 0.134* | -0.056* | -0.010* | 0.340 | -0.035* | 0.908 | -0.074* | -0.003* | -0.077* | -0.040* |
| Eudicots | 49 | -0.019* | -0.009* | 0.689 | 0.586 | 0.022* | 0.284 | -0.021* | 0.006* | -0.019* | 0.051* | 0.884 | 0.368 | -0.008* | 0.136 | 0.334 |
| **Metazoa** | 186 | 0.044 | 0.014* | 0.831 | 0.681 | -0.004* | 0.087 | 0.154 | 0.042 | -0.005* | -0.002* | 0.754 | 0.107 | 0.010* | 0.022 | 0.095 |
| Protostomia: | 78 | 0.025* | 0.025* | 0.867 | 0.679 | -0.010* | 0.034* | 0.167 | 0.059 | 0.003* | 0.011* | 0.772 | 0.047 | 0.001* | 0.013 | 0.059 |
| – Lophotrochozoa | 10 | 0.747 | 0.114* | 0.849 | 0.918 | 0.551 | 0.070* | 0.830 | 0.769 | 0.453 | 0.074* | 0.929 | 0.193* | 0.116* | -0.106* | 0.434 |
| – Arthropoda: | 61 | 0.019* | -0.014* | 0.858 | 0.723 | -0.0001* | 0.102 | 0.170 | 0.075 | 0.011* | -0.017* | 0.797 | 0.141 | -0.016* | 0.056 | 0.121 |
| – Diptera | 13 | -0.075* | 0.009* | 0.775 | 0.780 | -0.032* | -0.049* | -0.040* | -0.071* | 0.120* | -0.015* | 0.791 | -0.081* | -0.069* | -0.064* | -0.081* |
| – Hymenoptera | 18 | 0.533 | 0.007* | 0.864 | 0.284 | 0.365 | 0.166* | 0.629 | -0.053* | 0.430 | 0.117* | 0.321 | 0.156* | 0.015* | 0.072* | -0.056* |
| – Lepidoptera | 13 | -0.046* | -0.010* | 0.619 | 0.682 | 0.343 | -0.082* | 0.329 | -0.019* | 0.186* | -0.066* | 0.611 | 0.462 | -0.089* | -0.058* | 0.094* |
| Deuterostomia: | 101 | 0.027* | 0.049 | 0.841 | 0.730 | 0.028* | 0.205 | 0.101 | 0.006* | 0.063 | 0.056 | 0.743 | 0.248 | 0.261 | 0.163 | 0.104 |
| – Teleostei | 17 | 0.130* | 0.612 | 0.941 | 0.919 | 0.517 | 0.360 | 0.016* | 0.096* | 0.350 | 0.509 | 0.887 | 0.546 | 0.503 | 0.548 | 0.515 |
| – Amniota: | 72 | 0.0004* | -0.013* | 0.809 | 0.408 | -0.014* | 0.246 | 0.267 | 0.093 | 0.266 | 0.044 | 0.555 | 0.063 | 0.293 | 0.152 | 0.009* |
| – Reptiles | 11 | -0.009* | -0.111* | 0.295 | 0.211* | -0.052* | -0.091* | 0.455 | 0.380 | -0.095* | -0.107* | 0.695 | -0.085* | -0.084* | 0.117* | -0.108* |
| – Aves | 28 | 0.176 | 0.032* | 0.616 | 0.178 | 0.058* | 0.386 | 0.447 | 0.153 | 0.430 | 0.193 | 0.182 | -0.038* | -0.035* | 0.369 | 0.013* |
| – Mammalia | 33 | 0.025* | -0.018* | 0.782 | 0.629 | 0.017* | 0.150 | 0.267 | 0.055* | 0.218 | -0.006* | 0.742 | 0.004* | -0.032* | -0.024* | 0.085* |

Note: All genome-feature values were log10- transformed prior to analysis (see Methods). PGLS regressions are performed with estimated genome sizes and the “reference tree topology” (in Figure 1a). Symbology, **IS**: weighted-average intron size (nts), **IN**: total number of introns, **ID**: (absolute) intron density, **CDSI**: fraction (%) of intron-containing CDSs, **IC**: intronic content of the genome in Mbs (as based on assembled sizes), **IR**: fraction (%) of repetitive content within introns. Only $p > 0.05$ values are shown with asterisks, the remaining $p$ -values for $r^2$ have statistical significance: $< 0.001$ . The regressions performed with alternative tree topologies and estimated genome sizes are provided in Supplementary Table S13.

### Genome-wide intron features are decoupled among themselves at the local and broadest phylogenetic scales

The repeated decoupling of intron features from genome size evolution is also supported by two additional observations. First, the evolution of intron features appears to be differentially constrained by phylogeny rather than genome size throughout Eukarya. As shown in Figures 2-4 and also Supplementary Tables S14-S15, a homogeneous variation can be observed in the phylogenetic patterns of intron features across land plants, fungi and stramenophiles. This contrast the dramatic variation –at a much faster evolutionary pace– observed in the phylogenetic patterns of intron features within holozoans, chlorophytes, alveolates and amoebozoans. Second, the PGLS correlations in Tables 2 and 4 show that intron features associate weakly among themselves at the local and broadest phylogenetic scales (see also Supplementary Table S6-10 and S13). These associative correlations further support differing patterns between those features estimating intron abundance within and across genes (*i.e.*, intron density and the fraction of intron-containing genes, respectively) and those features measuring intron length and repeat composition. For instance, variation in intron size is weak but to a larger extent associated with the repetitive content within introns (*r*^2^ = 0.366, *logBF* = 588.2), rather than with changes in intron density (*r*^2^ = 0.025, *logBF* = 0.2), the number of introns per genome (*r*^2^ = 0.003, *logBF* = 1.6), or the fraction of intron-containing genes (*r*^2^ = 0.002, *logBF* = 1.6). Likewise, variation on intron density is weakly but to a larger extent associated with the number of introns per genome (*r*^2^ = 0.437, *logBF* = 161.5) and the fraction of intron-containing genes (*r*^2^ = 0.199, *logBF* = 152.0), rather than with changes in intron size, the repetitive content within introns (*r*^2^ = 0.210, *logBF* = 0.5) and the total number of CDS (*r*^2^ = 0.024, *logBF* = 0.074). As also observed in Table 4, the strenght of these associations fluctuates at local phylogenetic scales.

The evolutionary decoupling of intron features is also observed at the lineage and species levels (see summary statistics in Table 3 and Figures 3-4). For instance, Pezizomycotina and Eudicots show similar fractions of intron-containing genes on average (∼75%), although their corresponding avg. intron size (107.2 nts and 495.6 nts) and intron density (2.3 and 5) are different. Likewise, the small genomes of bees (288.7 Mbs) harbor shorter but more abundant introns within (6.2) and across (94.7%) genes than the larger genomes of butterflies (402.0 Mbs, 5.6 and 86.5% introns within and across genes, respectively). Strikingly, introns in Aves are slightly more abundant within (10.07) and across (92.9%) genes in comparison to mammals (9.3 and 86.8% introns within and across genes, respectively) (see Table 3 and Figure 4a). Even despite the fact that birds have undergone a reduction of their average intron (3,342 nts) and genome (1.2 Gbs) sizes, as observed here and elsewhere [76, 88, 89]. Consistent with this, birds and mammals show similar fractions of AS genes and have among the highest rates of AS events per gene [60]. Within fungi, we observe that introns within Basidiomycota are shorter (92.2 nts) but more abundant within (4.7) and across (81.8%) genes when compared to the larger (141.4 nts) but less abundant introns within (1.9) and across (53.4%) genes in Ascomycota (see Table 3 and Figure 3a). Indeed, an increase in the average rate of alternative splicing from 6% in Ascomycota to 8.6% in Basidiomycota has been previously reported [61]. Noteworthy, the large genome of the locust *L. migratoria* (6.5 Gbs) exhibits the lastest intron sizes of Eukarya [90]: ∼75% of its introns are between 5,000 and 50,000 nts. Surprisingly, the abundance of its introns within (5.73) and across (81.68%) genes is not higher with respect to other protostome clades. Additional examples are described in Supplementary Tables S14-S15.

**Figure 3.**
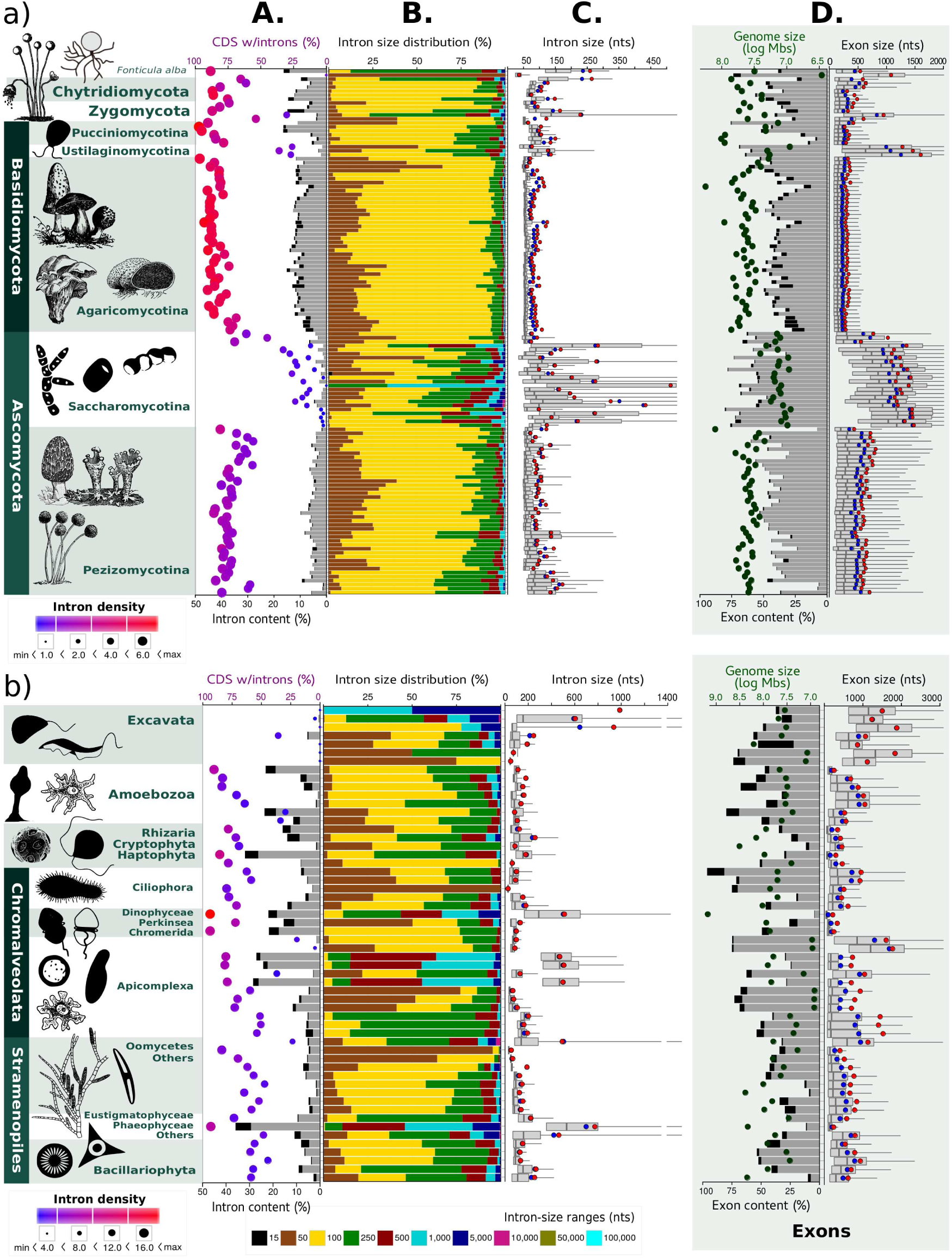
Phylogenetic distribution of intron and exon features across sequenced genomes in a) Fungi and b) Protists. **Panel A.** Distribution of intron content, density and fraction of protein-coding genes containing introns across species. Intron contents with unique and repetitive intronic sequences (based on estimated genome sizes) are shown in grey and black bars, respectively. Estimations based on the assembled genome sizes are plotted in Figures S7a-S10a. The information represented by the dots is two fold: 1) the fraction (%) of genes with introns is represented by the coordinate with respect to the top scale; 2) intron density is depicted by the size and the color of the dot, so that, a bigger dot with an intensified red color implays the presence of more introns per genes. **Panel B.** Intron size distribution within a genome is represented by the fraction (%) of introns from the total population binned in the following ranges: 15-50 nts (brown), 51-100 nts (yellow), <250 nts: introns presumably spliced by intron definition (green), >251 nts: introns presumably dismissed by exon definition. **Panel C.** Grey box-plots show the descriptors (quartiles, means and outlier-thresholds) for the intron length distribution in every genome; from right to left: Q1, median, Q3, upper-fence (line), standard average size (blue dot), and weigthed-average size (red dot). **Panel D.** Distribution of exon features. On the left side, bars show the exon content (bottom scale) with unique and repetitive exonic sequences in grey and black, respectively. Genome size (log Mbs) is represented by the coordinate with respect to the logarithmic top scale. On the right side, the exon size distribution within the genome is shown as described for introns in Panel C. Some upper-fences were cut to avoid a higher compactation of the data. The images for model species were kindly provided by silhouettesfree.com and ClipArt.com.

**Figure 4.**
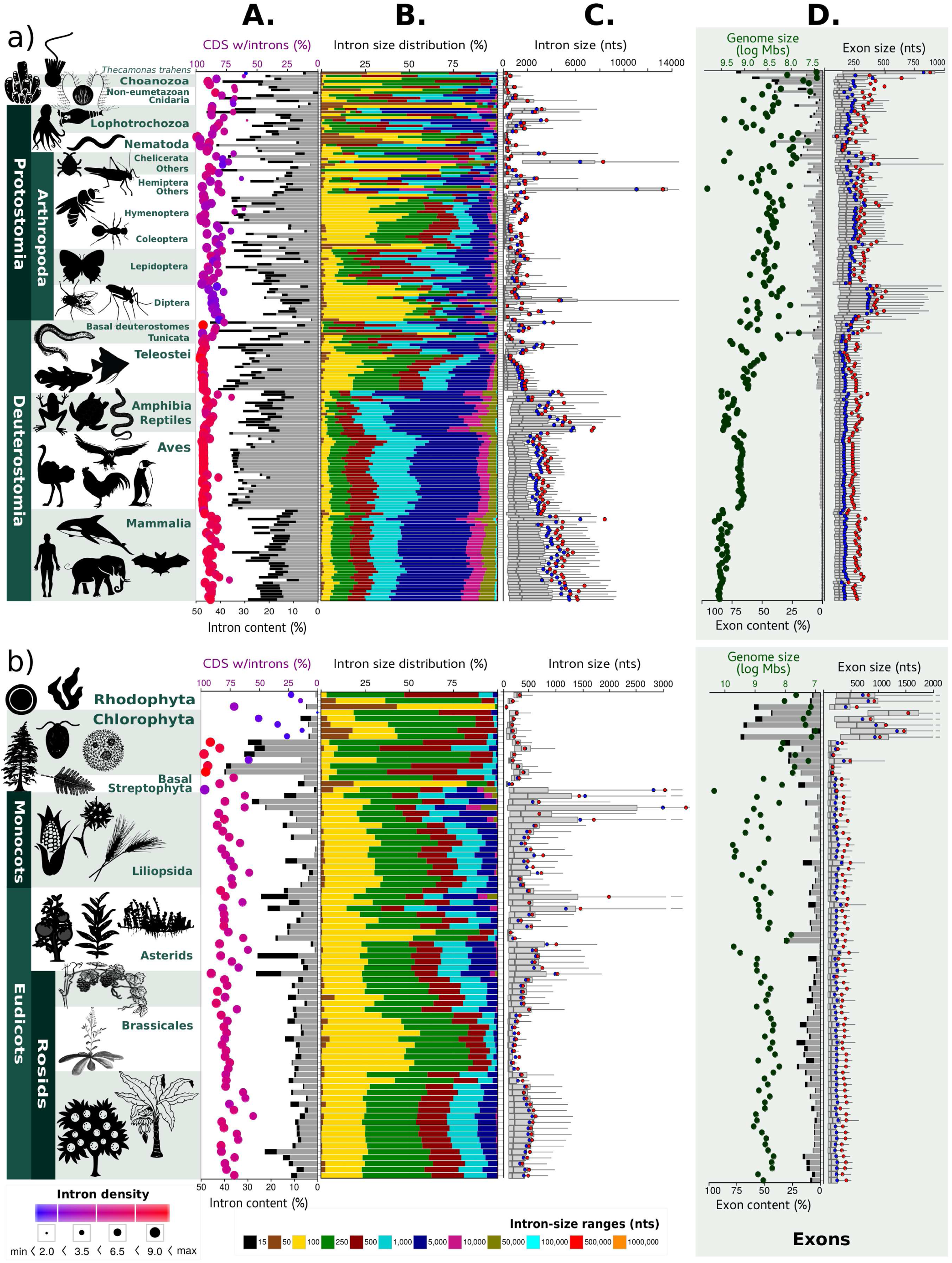
Phylogenetic distribution of intron and exon features across sequenced genomes in a) Metazoa and b) Archaeplastida. For panel description see Figure 3.

### Repeats differentially contribute to intron size and content across eukaryotes

We also investigated how significant is the contribution of repeats to the size and content of both introns and genomes. In partial agreement with previous studies [63, 64], we found that genome size is strongly associated at the broadest phylogenetic scale with its repetitive content (*r*^2^ = 0.723, *logBF* = 1273.3), but to a much lesser extent with the repetitive-intronic content (*r*^2^ = 0.485, *logBF* = 119.6) (see Table 2). As observed on Figure 2 and Table 3, repeats cover on average 13.6% of genome size in fungi, 41.6% in Viridiplantae, 26.5% in Metazoa, 17.7% in Choanozoa, 26.9% in Amoebozoa, 25.2% in Excavata, 26.7% in Stramenopiles, and 12.1% in Alveolata. When still observed at the supergroup level, the fraction of repeats covering the intronic genome content appears to mirror the previous trends: 8.8% in Fungi, 24.2% in Metazoa, 20.2% in Viridiplantae, 19.7% in Choanozoa, 29.6% in Amoebozoa, 13.0% in Excavata, 19.8% in Stramenopiles, and 14.6% in Alveolata. At the broadest phylogenetic scale, however, the repetitive content of the genome does not strongly correlate with intron size (*r*^2^ = 0.292, *logBF* = 56.4) or intron content (*r*^2^ = 0.314, *logBF* = 65.7), and it does not significantly associate either with intron density (*r*^2^ = 0.032, *logBF* = 3.1) or the fraction of intron-containing genes genes harboring introns (*r*^2^ = 0.018, *logBF* = 2.9) (see Table 2). These results show that the repetitive composition of introns is not strongly scaling with changes in genome size or the repetitive content of the genome at the broadest phylogenetic scale. Yet, fluctuations in the associations (from absent to strong) among intron features and the repetitive genome content are expected across lineages.

Figure 2 also shows whether or not the contribution of repeats to intron size is significant in every genome analyzed, according to the *p* < 0.001 obtained for the permutation tests performed on the *Jaccard index* (see Methods and Supplementary Table S16). In summary, we found no significant degree of nucleotide overlap between repeats and intronic sequences in most of the genomes analyzed in Fungi (71.0%), Viridiplantae (82.1%, with exceptions such as Prasinophytes), Aves, Oomycetes, Rhodophyta, and over half of the genomes within Mammalia. By contrast, the degree of overlap between repeats and intronic sequences was found statistically significant in most of the genomes analyzed within Metazoa (with notable exceptions), Amoebozoa (71.4%), and Alveolata (80%) (see Supplementary Table S16). Examples of species and clades with considerable high repetitive-intron contents (25-50%) and significant nucleotide overlaps between repeats and introns include: the haptophyta *Emiliania huxleyi*, the sea sponge *Amphimedon queenslandica*, Dictyostelia, Prasinophytes, Cnidaria, protostomates such as Lophotrochozoa, Diptera and Lepidoptera, as well as vertebrates such as Teleosts, Anura, Turtles, Crocodylia and Mammalia. With exception of Prasinophytes, high repetitive-genome contents (25-40%) are also observed in the previous lineages. Conversely, an apparent concerted reduction in the contribution of repeats to genome size (6-15%) and intron content (3-14%) –also supported by a non-significant overlap between repeats and intronic sequences– is observed in most species within Ascomycota, Nematoda, Hymenoptera, Aves, and in *Trichoplax adhaerens*. This particular pattern requires further research. Additional examples with statistic estimators are revisited in Supplementary Tables S14-15, S18.

### Genome-wide exon features are also weakly associated with genome size, but evolving steadier than intron features

We further estimated the phylogenetic patterns of exons features across eukaryotes to contrast those observed among intron features. As shown in Figures 2-4 and Table 5, around half of the genome size is covered by exonic sequences in Fungi (44.9%), Amoebozoa (53.8%), Alveolata (51.9%), Prasinophytes (81.5%) and Excavata (53.1%). In contrast, low fractions of exon content are observed in the genomes of land plants (13.8%) and metazoans (5.2%), with pinus and mammalian genomes harboring barely 1% of protein-coding nucleotides. As with intron features, we found that the variation estimated for exon features is weakly associated to the variation observed in eukaryotic genome sizes at the broadest phylogenetic scale. As observed in Table 2 and Supplementary Tables S6-S10, genome size is positive but to a lesser extent associated with: exon content (*r*^2^ = 0.262, *logBF* = 6.2), the total number of both exons (*r*^2^ = 0.316, *logBF* = 11.3) and CDS (*r*^2^ = 0.328, *logBF* = 20.3), as well as the average length of CDS (*r*^2^ = 0.263, *logBF* = 43.8). These estimations are in agreement with previous studies [64, 66]. However, none or no simple associations were found for genome size against exon density (*r*^2^ = 0.054, *logBF* = 0.05), exon size (*r*^2^ = 0.096, *slope* = -0.114, *logBF* = 0.2), and the repetitive-exon content (*r*^2^ = 0.135, *logBF* = 0.9).

**Table 5.** Summary statistics, mean value and (coefficient of variation), of several protein-coding features for different species datasets.

| Genome features<br>Taxon | *n* | Genome size<br>[Mbs] | CDS:<br>number | size<br>[nts] | content<br>[%] | EXONS:<br>size<br>[nts] | content<br>[%] | repeats<br>[%] | number |
| --- | --- | --- | --- | --- | --- | --- | --- | --- | --- |
| **Protists** | 66 | 83.6 (2.23) | 15,354 (0.90) | 2,071 (0.66) | 52.2 (0.39) | 804.3 (0.51) | 43.9 {44.3}(0.47) | 11.8 (0.78) | 58,467 (1.58) |
| Stramenopiles | 17 | 86.0 (0.82) | 16,190 (0.43) | 1,803 (0.66) | 41.4 (0.38) | 687.5 (0.30) | 32.9 {33.4}(0.44) | 13.0 (0.62) | 45,396 (0.62) |
| Alveolata | 20 | 116.1 (2.82) | 13,546 (0.90) | 2,671 (0.75) | 58.0 (0.37) | 880.7 (0.52) | 51.9 {51.7}(0.46) | 8.1 (0.76) | 82,731 (1.90) |
| **Fungi** | 131 | 38.2 (0.61) | 11,779 (0.44) | 1,500 (0.13) | 51.9 (0.27) | 626.3 (0.59) | 44.9 {45.1}(0.32) | 6.4 (1.16) | 44,567 (0.70) |
| Basidiomycota | 53 | 47.3 (0.56) | 14,626 (0.38) | 1,545 (0.16) | 51.9 (0.25) | 373.9 (0.72) | 41.5 {41.0}(0.30) | 10.4 (0.82) | 72,171 (0.37) |
| Ascomycota: | 66 | 30.8 (0.59) | 9,464 (0.38) | 1,467 (0.08) | 52.2 (0.29) | 839.8 (0.38) | 48.2 {48.7}(0.33) | 2.3 (0.94) | 22,788 (0.59) |
| – Pezizomycotina | 42 | 40.6 (0.39) | 11,519 (0.25) | 1,488 (0.10) | 45.4 (0.29) | 623.9 (0.14) | 40.3 {40.8}(0.30) | 2.2 (1.10) | 31,274 (0.28) |
| – Saccharomycotina | 22 | 13.7 (0.29) | 5,848 (0.14) | 1,432 (0.04) | 63.9 (0.18) | 1,246.8 (0.16) | 62.1 {62.8}(0.19) | 2.5 (0.70) | 7,520 (0.47) |
| **Chlorophyta** | 10 | 49.3 (0.91) | 10,349 (0.40) | 2,603 (0.52) | 67.9 (0.36) | 610.7 (0.78) | 47.7 {45.5}(0.72) | 7.9 (0.63) | 56,632 (0.93) |
| **Streptophyta** | 68 | 1,065.8 (2.54) | 35,359 (0.51) | 3,053 (0.56) | 27.3 (0.75) | 274.7 (0.46) | 13.8 {15.1}(1.34) | 16.4 (0.82) | 171,871 (0.55) |
| Monocots | 15 | 1,809.8 (0.96) | 37,055 (0.40) | 3,619 (0.59) | 15.0 (0.78) | 385.3 (0.17) | 5.2 {6.4}(0.91) | 13.3 (0.82) | 171,992 (0.30) |
| Eudicots | 49 | 641.1 (1.23) | 40,012 (0.43) | 2,859 (0.49) | 23.7 (0.42) | 377.0 (0.11) | 10.1 {12.3}(0.63) | 19.5 (0.74) | 197,128 (0.49) |
| **Metazoa** | 186 | 1,261.0 (0.93) | 18,413 (0.34) | 17,906 (0.77) | 30.6 (0.40) | 287.7 (0.24) | 5.2 {5.4}(1.13) | 6.9 (0.89) | 142,824 (0.43) |
| Protostomia: | 78 | 517.9 (1.58) | 17,266 (0.44) | 7,223 (0.94) | 30.8 (0.44) | 322.3 (0.24) | 7.8 {8.3}(0.72) | 7.2 (0.80) | 93,884 (0.53) |
| –Lophotrochozoa | 10 | 765.6 (0.97) | 26,623 (0.45) | 8,316 (0.63) | 29.6 (0.37) | 308.3 (0.11) | 6.8 {7.7}(0.94) | 15.2 (0.38) | 153,645 (0.39) |
| –Arthropoda: | 61 | 523.3 (1.66) | 15,608 (0.36) | 7,534 (0.96) | 29.2 (0.46) | 340.3 (0.20) | 7.0 {7.5}(0.66) | 5.9 (0.80) | 81,455 (0.44) |
| – Diptera | 13 | 410.1 (0.66) | 13,435 (0.22) | 5,975 (0.64) | 22.4 (0.40) | 416.7 (0.17) | 6.9 {7.9}(0.71) | 4.4 (0.50) | 54,745 (0.33) |
| – Hymenoptera | 18 | 278.6 (0.27) | 13,584 (0.25) | 7,138 (0.48) | 33.3 (0.36) | 328.8 (0.16) | 7.4 {7.9}(0.31) | 5.2 (0.76) | 74,930 (0.13) |
| – Lepidoptera | 13 | 402.0 (0.37) | 16,232 (0.25) | 7,279 (0.30) | 32.6 (0.51) | 300.8 (0.14) | 5.2 {5.2}(0.38) | 4.0 (0.63) | 96,678 (0.39) |
| Deuterostomia: | 101 | 1,898.6 (0.55) | 19,010 (0.24) | 27,113 (0.42) | 29.7 (0.28) | 254.4 (0.15) | 2.5 {2.8}(1.30) | 5.7 (0.73) | 182,585 (0.21) |
| –Teleostei | 17 | 871.6 (0.39) | 22,109 (0.10) | 14,118 (0.34) | 36.5 (0.16) | 235.8 (0.14) | 4.7 {5.2}(0.35) | 5.1 (0.51) | 235,557 (0.12) |
| –Amniota: | 72 | 2,234.2 (0.43) | 17,793 (0.16) | 32,314 (0.25) | 28.0 (0.25) | 255.4 (0.12) | 1.5 {1.6}(0.37) | 5.0 (0.67) | 173,398 (0.14) |
| – Reptiles | 11 | 2,188.3 (0.24) | 18,194 (0.13) | 33,886 (0.29) | 28.0 (0.16) | 272.9 (0.08) | 1.4 {1.4}(0.30) | 8.4 (0.35) | 163,293 (0.16) |
| – Aves | 28 | 1,231.1 (0.11) | 15,031 (0.09) | 27,634 (0.16) | 33.7 (0.15) | 232.4 (0.05) | 2.1 {2.2}(0.11) | 2.2 (0.38) | 157,084 (0.06) |
| – Mammalia | 33 | 3,100.6 (0.16) | 20,003 (0.09) | 35,762 (0.23) | 23.2 (0.23) | 269.0 (0.12) | 1.0 {1.2}(0.18) | 6.3 (0.47) | 190,608 (0.11) |

Note: Genome contents provided in % or Mbs are calculated from assembled genome sizes. Unique or non-repetitive exon contents are denoted with {}. Summary statistics for additional clades and for genome contents based on estimated genome sizes are provided in Supplementary Tables S14-S15.

Contrasting the intron size’s patterns, the distribution of exon size is tighter across eukaryotes (see Tables 3 and 4). We found that between 50% and 75% of the exon population within a genome has a narrow and small length below 250 nts across plants, green algae, Choanozoa, basiodiomycetes, and Metazoa. Accordingly, we observed that exon length distributions are less-skewed in most eukaryotic clades (Figures 3D and 4D) when compared to intron length distributions (Figures 3C and 4C). Nevertheless, large exon lengths are observed in Rhizaria (407.2 nts), Amoebozoa (600.7 nts), Stramenopiles (687.5 nts), Alveolata (880.7), Excavata (1,357.3 nts), Prasinophytes (985.9 nts), Rhodophyta (870.9), Ustilaginomycotina (1,107.1 nts) and Ascomycota (721.6 nts), particularly in Saccharomycetes (1,167.23 nts) (see Figures 3D-4D and Supplementary Figures S7-S10). As observed in Table 2, furthermore, two different correlative associations were found for exon size at the broadest phylogenetic scale. On the one hand, exon size significantly decreases as the density and total number of exons in the genome increases (*r*^2^ = 0.731, *slope* = -0.880, *logBF* = 97.5 and *r*^2^ = 0.517, *slope* = -1.305, *logBF* = 10.5, respectively). On the other hand, none or no simple associations were found between exon size and the number (*r*^2^ = 0.097, *logBF* = 0.6) or length of CDS (*r*^2^ = 0.050, *logBF* = -0.01). These results are in overall accordance with previous estimations [66, 91, 92].

As observed in Table 5 and Figure 2, the fraction of repeats covering the exonic genome content across eukaryotes is considerably small (between 2% and 8%), in comparison to introns. Although larger fractions are observed in some species within land plants (23.05%), Lophotrochozoa (15.2%), Cnidaria (26.3%), Basiodiomycota (10.4%), Excavata (17.5%) Amoebozoa (16.2%) and Stramenophiles (13.0%). In agreement with these observations, the average length of exons is not significantly associated to its repeat content (*r*^2^ = 0.034, *slope* = -0.564, *logBF* = 2.3) or the genome repeat content (*r*^2^ = 0.094, *slope* = -1.269, *logBF* = 0.1). Based on the results from the *Jaccard index*’s permutation tests (see Figure 2), we found no significant degree of overlap between repeats and exonic sequences across most of the eukaryotic genomes analyzed here: Amoebozoa (100%), Choanozoa (100%), Metazoa (97.1%), Fungi (92.4%), Excavata (92.4%), Viridiplantae (86.5%), Stramenophiles (82.4%) and Alveolata (80%). The noteworthy exceptions can be observed in Figure 2 and Supplementary Table S16.

### Intron-richness is robustly associated to complex multicellular organisms and their closest ancestral relatives

We further investigated the relationship between genome-wide intron features and multicellular complexity. As described in Appendix 1, we developed four criteria and three definitions to distinguish the species in our dataset as: complex multicellular (CMOs: 288), simple multicellular (SMOs: 96) or unicellular (77) organisms (see Supplementary Table S2). Accordingly, CMOs are defined here as those *organisms exhibiting an irreversible transition in individuality produced by tissue-based body plans, through the developmental commitment of multiple and different cell types originated from a common cell-line ancestor*. We then used Principal Component Analyses (PCAs) with direct comparative data (compPCA) and phylogenetically independent contrasts (phyloPCA) to investigate how seven intron features are covarying among themselves (Figure 5b), with other eight genome features (Figure 5e), with complex multicellularity (Figure 5a,d), and with genome size (Figure 5c,f). Noteworthy, the findings that are going to be described next, also hold for both the com-PCA analysis and the phyloPCAs performed with different tree topologies and assembled genome sizes (see Supplementary Figures S11-S15 and Tables S18-S20). As observed in Figure 5, high intron-richness does segregate CMOs (in red) from both SMOs (in blue) and unicellular organisms (in yellow), by clustering through genome-wide intron features exclusively and in conjunction with other genome features. The first two principal components (PC1 and PC2) from both phyloPCA analyses capture most of the variances in the data: 87.16% with 7 intron features and 64.6% with 15 variables. However, the association of the variables on each principal component is different in both phyloPCAs (see Table 6), as described next.

**Table 6.** Contribution (%) of 7 intron features, 15 genomic traits and cellular complexity to the first seven principal components as estimated with the phylogenetic Principal Component Analyses (phyloPCA).

| *Intron features only* | **PC1** | **PC2** | **PC3** | **PC4** | **PC5** | **PC6** | **PC7** |
| --- | --- | --- | --- | --- | --- | --- | --- |
| Cumulative % of var | **68.09** | **19.07** | **7.04** | 3.22 | 2.2 | 0.24 | 0.15 |
| Intron number | **16.935** | 10.744 | 0.666 | 4.684 | 18.121 | 41.191 | 7.659 |
| CDS with introns (%) | 11.568 | **19.174** | **20.104** | 25.391 | 23.682 | 0.026 | 0.055 |
| Intron density | 11.763 | 6.245 | **67.774** | 0.539 | 13.618 | 0.059 | 0.001 |
| Intron weighted-mean size | 4.864 | **54.072** | 0.803 | 14.782 | 3.767 | 16.261 | 5.451 |
| Intron content | **19.986** | 2.249 | 0.062 | 0.543 | 5.538 | 0.042 | 71.58 |
| Unique intron genome | **19.665** | 0.775 | 0.218 | 4.937 | 20.867 | 39.104 | 14.434 |
| Intron repeats | **15.219** | 6.741 | 10.373 | 49.123 | 14.406 | 3.317 | 0.82 |
| *15 traits* | **PC1** | **PC2** | **PC3** | **PC4** | **PC5** | **PC6** | **PC7** |
| Cumulative % of var | **50.18** | **14.39** | **12.28** | 6.2 | 5.54 | 4.27 | 3.2 |
| Cellularity type | 1.250 | 0 | 0.020 | 95.603 | 0.368 | 0.600 | 1.848 |
| Number CDS | 6.842 | **12.412** | 6.212 | 0.001 | 0.001 | 1.717 | 2.969 |
| CDS with introns (%) | 3.316 | **15.560** | 7.007 | 0.281 | 5.366 | 12.077 | 30.978 |
| Genome size | **9.195** | 0.980 | 9.305 | 0.030 | 0.472 | 8.715 | 6.303 |
| Repeat genome content | 7.262 | 1.852 | 10.440 | 0.138 | 12.259 | 5.600 | 0.684 |
| Unique nc-genome content | **9.476** | 0.009 | 6.467 | 0.012 | 2.063 | 9.464 | 4.721 |
| Intron content | **11.637** | 2.943 | 0.007 | 0.438 | 0.902 | 6.417 | 0.004 |
| Unique intron genome | **10.966** | 4.039 | 0.143 | 0.346 | 3.140 | 4.793 | 0.046 |
| Intron repeat content | **10.253** | 0.312 | 1.742 | 0.387 | 8.792 | 7.684 | 0.439 |
| Intron density | 4.597 | **15.291** | 5.384 | 0.290 | 1.476 | 5.013 | 26.913 |
| Intron weighted-mean size | 3.757 | 0.075 | **27.306** | 1.682 | 7.321 | 10.892 | 9.427 |
| Exon average size | 4.796 | **14.971** | 4.033 | 0.067 | 2.436 | 24.303 | 1.409 |
| Exon content | 6.528 | **12.239** | 9.952 | 0.381 | 3.547 | 0.294 | 0.356 |
| Unique exon content | 6.155 | 8.347 | 10.293 | 0.156 | 15.378 | 0.197 | 0.058 |
| Exon repeat content | 3.971 | **10.971** | 1.689 | 0.188 | 36.480 | 2.236 | 13.845 |

Note: Since all genome-feature values were log10- transformed prior to analysis, phyloPCAs were performed with 457 species (see Methods). Only results from the phyloPCAs performed with the reference tree topology (Figure 1a) and genome features based on assembled genome sizes are shown here. Results for the phyloPCAs performed with other tree topologies and estimated genome sizes, as well as from the comparative PCA, are provided in Supplementary Tables S13-S14. The noticeable contribution of some variables for the first three PCA components is highlighted with bold font.

**Figure 5.**
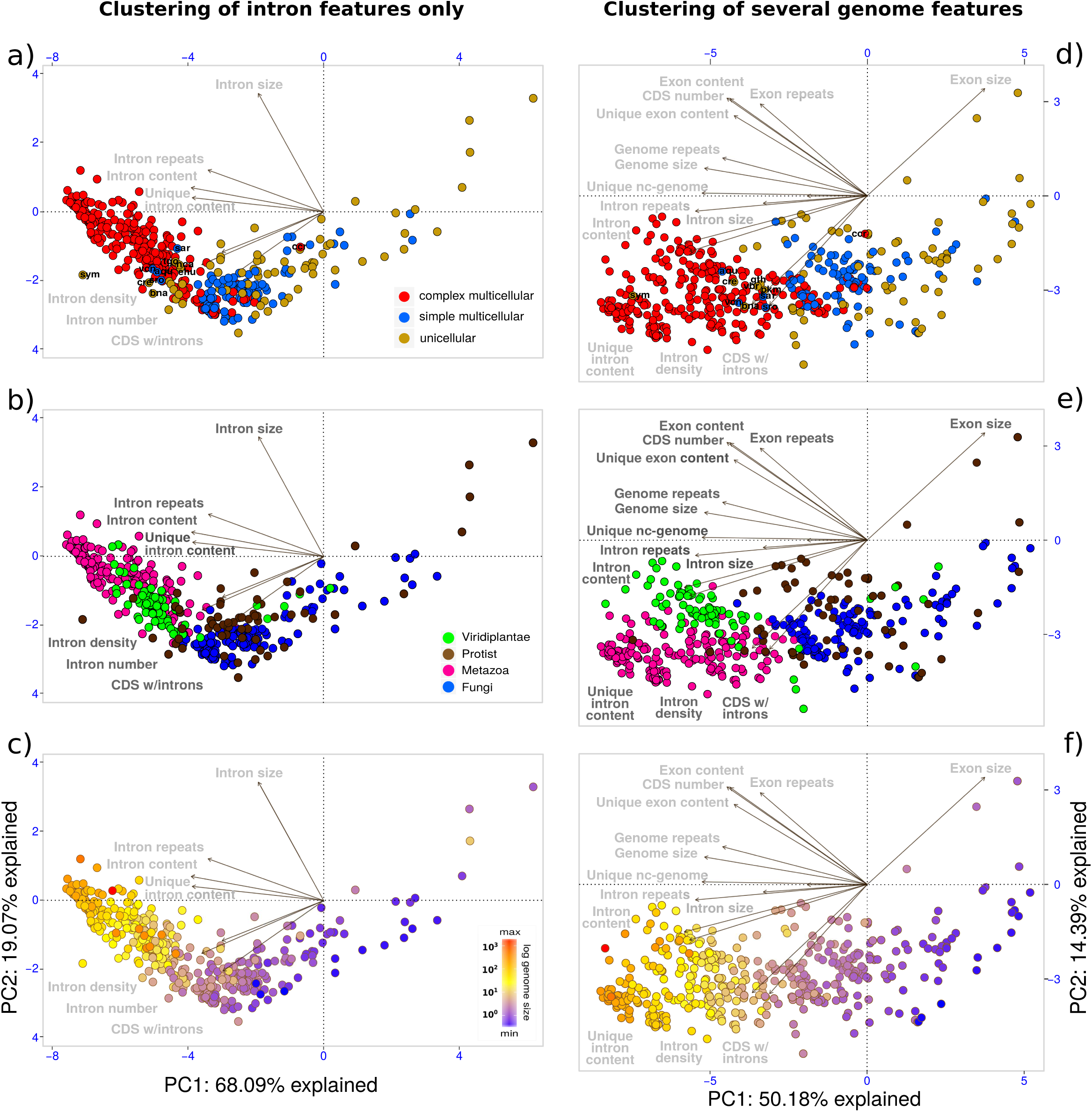
Phylogenetic principal component analysis (phyloPCA) of introns, other genome features and multicellular complexity. The phyloPCAs were performed with the reference tree topology (Figure 1a). The biplots depict the two first components (PC1 and PC2) inferred from the phyloPCA of 7 intron features (a-c), 14 genomic traits (d-f) and the organism complexity (a and d) estimated for 457 sequenced eukaryotes. In plots a) and d), the species are color-coded according to their organismal complexity (see Appendix 1): unicellular (yellow), simple multicellular (blue), complex multicellular (red). In plots b) and e), species are color-coded by major eukaryotic supergroups: Fungi (blue), Metazoa (pink), Viridiplantae (green), and “protists” (brown). In plots c) and f), species are color-coded by their log10-transformed genome sizes. Species are clustered in the phyloPCAs according to their dispersion along PC1 and PC2, with a confidence limit of 0.95. The dark lines radiating from (0,0) represent each variable included in the analysis; the direction of a line represents the highest correlation coefficient between the scores of the principal components and the variable, while its length is proportional to the strength of this correlation. Noteworthy, the Cronbach’s alpha coefficients obtained for the sets of 7 and 15 variables (0.88 and 0.95, respectively) indicate high internal consistency among the variables to measure the same underlying concept through phyloPCA analyses (see Methods). PhyloPCA analyses performed with alternative tree topologies and assembled genome sizes are provided in Supplementary Tables S16-S20 and Figures S11-S15.

By analyzing the phyloPCA of intron features, we first observe that high intron-richness clusters CMOs together (Figure 5a), regardless of the lineage they belong to (Figure 5b) or the wide dispersion of their genome sizes (Figure 5c). We further observe that the content, repeat composition and number of introns are mainly contributing to PC1 (71.82%), while PC2 has a major contribution (73.24%) from the fraction of intron-containing genes, intron size and density (see Table 6). In the broarder phyloPCA analysis that includes additional genome features, we also found that intron features are leading the clustering of CMOs, while other genome features are clearly setting apart the major eukaryotic supergroups: metazoans (in pink), plants (in green), fungi (in blue) and protists (in brown). Consequently, the association between high intron-richness and CMOs is found to be in part a byproduct of other genomic features constraining the development of multicellularity in every supergroup. For instance, PC1 clearly captures the *ncDNA complexity* of the genomes, while PC2 captures their *protein-coding complexity*. Accordingly, PC1 has a major contribution (51.5%) from intron content, unique intron content, unique ncDNA, genome size and genome repeat content (see Table 6). Notably, the scaling distribution of genome sizes along PC1 in Figure 5f endorses the strong correlative association found between ncDNA and genome size at the broadest phylogenetic scale (*r*^2^ = 0.859, see Table 2). Conversely, exon features (size, content, repeat composition), the number of CDS, the fraction of intron-containing genes and intron density contribute mostly to PC2 (89.8%). Hence, intron-richness in land plants is mainly associated with the high contribution of repeats to the genome, while intron-richness in metazoans is mostly related to the non-repetitive ncDNA fraction of the genome. By contrast, most unicellular and SM species (in fungi and protists) are strongly associated with several exon features, particularly with exon size and the number of CDS.

The previous patterns strongly suggest that high intron-richness is a robust genomic fingerprint of both CMOs and their closest ancestral relatives. For instance, ancestral relatives clustered among CMOs include: the choanoflagellates *Salpingoeca rosetta* and *Sphaeroforma arctica*, the sponge *A. queenslandica*, the slime mold *Fonticula alba*, the green alga *Volvox carteri*, and other chlorophytes (see Figure 5a,d). Nevertheless, there are some exceptions to this trend. First, there are also few exceptional intron-rich unicellular organisms clustered among CMOs, with no evidence of multicellularity (either simple or complex): the dinoflagellate *Symbiodinium minutum*, the chlorarachniophyte alga *B. natans*, the green alga *Chlamydomonas reinhardtii* and the apicomplexans *Toxoplasma gondii, Neospora caninum* and *Hammondia hammondi*. The high intron-richness of these unicellular species –comparable to the one observed in vertebrates– has been already acknowledged, and is suggested to have a role in the development of their complex life cycles [93–96]. Second, we observed a very few intron-poor CMOs, such as the red algae *Chondrus crispus* and the highly derived cnidarian parasite *T. kitauei*, that have undergone massive loss of introns and genome reduction due to extreme lifestyle conditions [44, 97]. Until more genomes of multicellular red algae are sequenced, however, it would be unclear to know whether the relative intron-poorness observed in *C. crispus* is an ancestral constraint or a derived (exceptional) trait [44]. Despite the presence of few intron-poor CMOs, the statistical robustness of the association between complex multicellularity (CM) and intron-richness is provided by most of the 288 intron-rich species classified as CMOs in this study, rather than by the five so far (out of the six) independent instances where CM has evolved. This is because intron loss overcomes intron gain across eukaryotes [1, 3, 4, 7] and intron-richness does not necessarily depend on genome size (this study). Therefore, there are no reasons to expect that high intron-richness has been maintained from intron-rich ancestors nor that it is only present in CMOs with large genome sizes. Yet, the major inconsistency found on both phyloPCA analyses (Figure 5) is the absence of a clear separation between some fungal SMOs and CMOs. This can be explained by the difficulty to determine the precise multicellular lifestyle of several fungi as simple or complex, owing to the lack of detailed life-cycle descriptions and the presence of “fuzzy” fruiting body development that challenge the evaluation of the criteria 3 and 4 in our definitions (see Appendix 1). This is an issue not only faced here, but also discussed elsewhere [98–101], and thus require further research.

## DISCUSSION

### Spliceosomal introns form a dynamically evolving ncDNA class, most likely under the influence of diverse life-history factors and evolutionary forces

Consistent with previous results [45, 66, 92] and with some noteworthy exceptions, our findings show that exons within protein-coding genes have remained within a narrow average size between 150 and 300 nts in Metazoa, Choanozoa, Viridiplantae and Basidiomycota. Hence, we observe that significant modifications in the structure of protein-coding genes in these lineages are basically a consequence of changes in the length and abundance of introns. Unexpectedly, these and other genome-wide features of introns are found to be repeatedly decoupled among themselves and from genome size evolution throughout Eukarya. Three findings support this observation. First, the strength of the associations among genome size and genome-wide intron features is different at the lineage-specific level, as consistent with other studies [76, 81, 102, 103]. Second, the features estimating the length and abundance of introns in a genome are weakly associated among themselves at the local and broadest phylogenetic scales. This explains the heterogeneous and contrasting patterns of intron evolution reported in the literature [2, 7, 55, 70, 76, 81]. As a consequence of the previous findings, changes of intron content cannot fully account for the large variations of eukaryotic genome sizes (in agreement with [64]), nor be strongly associated to the variation of one particular intron feature. Third, the repetitive composition of introns is not necessarily scaling with changes in genome size or the repeat content of the genome. Indeed, introns are found to be far from representing repetitive sequences in several lineages. As argued below, these results do not contradict –and even endorse in several cases– the contribution that different repeat classes have to either the origins [9, 11, 12] or length extension [73–76, 103] of introns at particular lineages. Therefore, our findings collectively unveil spliceosomal introns as a dynamically evolving ncDNA class.

Can a major mechanism (adaptive and non-adaptive) offer a unifying explanation to the highly heterogeneous patterns of intron evolution and genome complexity? Our findings suggest that this is highly unlikely. None of our correlative analyses imply causation nor offer evidence for the evolutionary mechanisms explaning the phylogenetic patterns reported here. Yet, strong (linear) correlations among several measures of genome complexity haven been often provided as evidence, by some evolutionary models [62, 65], to imply a concerted evolution among genome size, ncDNA content and intron-richness across eukaryotes (also discussed in [37, 67]). We observe here that genome size, its overall ncDNA and repetitive contents are indeed strongly associated at the large evolutionary scale (see Table 2). However, we also demonstrate that changes in the variation of some intron features (such as size and repeat composition) are only weakly, while other features measuring intron abundance are not, scaling with changes in genome size at the broadest phylogenetic scale. Our findings are thus in clear disagreement with previous estimations claiming the opposite [62, 63, 65, 66, 71]. Moreover, our results show that the genome-wide features determining length and abundance of introns across protein-coding genes are largely evolving independently throughout Eukarya. Our results are thus inconsistent with both a particular intron feature as key determinant of eukaryotic gene architecture, as well as a major mechanism (adaptive or non-adaptive) underlying a concerted effect between genome size and intron-richness over a large phylogenetic scale. Instead, the repeated decoupling among intron features themselves and with genome size strongly suggests that the major genome-wide features of introns from coding genes are evolving under the influence (direct or indirect) of either different or several life-history factors and evolutionary forces.

For instance, non-adaptive mechanisms such as the longterm evolutionary dynamics of repeats –which depend on factors like methylation propensity, RNAi-mediated interference and mating system– are found to determine concerted changes of genome and intron size in certain lineages [67, 73]. Examples observed here and elsewhere include some species within red algae and plants [44, 74, 103–105], insects [90, 106], fish [75] and birds [76, 88, 89]. Other studies suggest that intron density is determined by mechanistic factors such as nonhomologous end-joining (NHEJ) of DNA segments, reverse transcriptase and transposition activities [7, 70]. Also, introner-like elements greatly contribute to episodic intron gains in some algae [9, 12] and fungi [11]. And recently, rates of spontaneous intron-creating and -deleting mutations were found to shape the intron-exon structures of several distantly related species [107]. Under adaptive forces, variations of intron size and density across several eukaryotic lineages have been associated with the action of natural selection: (a) to conserve regulatory binding sites [17, 20, 21, 23] and regulatory ncRNAs [108–110]; (b) to promote the creation of new exons in vertebrates [49, 111]; (c) to reduce splice error rates [68] and protect against transcription-associated genetic instability [52]; (d) to favor co-transcriptional splicing and nucleocytoplasmic export of highly expressed and rapidly regulated cell-cycle genes [34, 37, 112]; (e) to reduce the metabolic costs associated with either powered flight in birds [76, 88] or environmental changes of habitats in teleosts [113].

Consequently, our findings also endorse concerns [67, 114, 115] regarding how much of the content, variation and complexity of intron-richness (along with other ncDNA classes) in Eukarya can be explained by the strong action of *N_e_* and genetic drift over a large evolutionary scale, as the “mutational-hazard” (MH) model states [62, 63, 80, 116]. In addition to the inconsistencies discussed previously, other studies [67, 68, 86] were not able to find statistically significant associations among *N_e_µ* and several intron and genome features after removing the phylogenetic signal from the dataset of Lynch and Conery [62]. Another studies have also shown that *N_e_* cannot exclusively or even largely explain major changes of intron and genome sizes in lineages within amniotes [76], insects [117], ascomycetes [81], and plants [67, 86]. Furthermore, reliable estimations of *N_e_* (such as *N_e_µ* and *K_a_*/*K_s_*) are still a matter of debate [67, 118], since they do not correlate well [68, 119] and are affected by several life-history traits [114, 120]. It is important to note that our findings do not dismiss the impact that *N_e_* and genetic drift has on the accumulation of ncDNA in eukaryotes. Rather, they argue that the population genetic settings suggested by the MH model are most likely to be dominant at the local phylogenetic scale, or in particular intron features from coding genes, or during recent founder events, or over introns located at non-coding regions.

### The robustness of systematic and phylogenetically controlled analyses

The results summarized above are based on a phylogenetic controlled framework over the largest and most diverse dataset of eukaryotic complete genomes to date. A major concern is, however, that phylogenetic uncertainty might affect considerably any phylogenetically controlled analysis [67, 82]. Here, we recapitulated no significant changes in our results after taking into account significant and numerous phylogenetic disagreements from four tree topologies estimated for the 461 species analyzed in this study. We could argue that the literature-based tree might reflect better the community consensus about the evolutionary history of these eukaryotes, since it is consistent with the *Open Tree of Life* [85] and with the species phylogenies reported with the complete genomes. However, the backbone of the tree of eukaryotes is still subject to deep rearrangements and competing hypothesis [84, 85]. Therefore, we can never exclude the possibility that a suggested tree is free of errors nor the existence of a better phylogenetic representation. Here, we have demonstrated that our phylogenetic controlled analyses are strongly robust to: (a) uncertainties about ancestral branches, such as Parahoxozoa in Metazoa [121, 122] or Excavata in Eukarya [84]; (b) discrepancies about the phylogenetic position of particular clades among their “peers”, for instance, Microsporia within Fungi [50] and Rhizaria within SAR [84]; (c) uncertainties in poorly resolved branches, such as Arthropoda and Rhizaria [123, 124]; as well as (d) common tree reconstruction problems, such as the presence of hard polytomies and long branch attraction of species.

In fact, the most significant discrepancies of our regressions from previous estimations over a large evolutionary scale are caused by the absence of correction for phylogenetic signal, rather than by the accuracy of the topological information used to account for it. As also shown by Whitney *et al.* [67] and Wu and Hurst [68], uncorrected phylogenetic dependencies among species (which assumes a star polytomy) lead to a much stronger correlation signal, as those strong correlations reported by previous studies [62, 65, 66]. We further showed that the correlation signal can also be affected considerably under controlled phylogenetic analyses, as some of those correlative associations estimated recently [64, 68]. Some of the factors analyzed here that lead to such biases include: low phylogenetic diversity, very small species datasets (<100 species) attempting to represent the current diversity of the sequenced eukaryotes, and the lack of systematic estimations of genome traits. These problems are particularly found at the clade-specific level (as also reported in [76, 81]), and in biased estimations of intron features (*e.g.*, density and genomic content) that already incorporate changes of genome size or on the number of protein-coding genes (see Appendix 2). We also demonstrated that our results are robust, not only to a reduced number of species (as long as the phylogenetic diversity of the dataset is maintained), but also to different sources of genome size estimates.

Yet, two additional factors might challenge the findings of this study. First, a limited access to both larger genomes (>5 Gbps) and genomes from deep branches of the eukaryotic tree precludes us from evaluating eventual biases on phylogenetic and “C-value” diversity so far. Second, substantial errors in genome assembly and/or annotation of protein-coding genes might considerably affect estimations of genome-based features. For instance, under- and over-estimation of genomic features were recently found on the genome projects of *Branchiostoma floridae* and *Hydra magnipapillata*, respectively, after RNA-seq-based re-annotations were performed in seven holozoans [66]. It is unclear, however, which genome features are the most affected by these biases, because the method employed in such study cannot distinguish introns and exons of coding genes from non-coding genes, such as pseudogenes and ncRNAs. This distinction is relevant in our study, since coding genes have a different copy number and experience different selective pressure than non-coding genes [125–127]. Moreover, annotation of protein-coding genes is becoming increasingly reliable owing to the incorporation of unbiased RNA-seq data, as most of our annotations are supported by. Through the filtering approaches of `GenomeContent`, we further found that very small introns and exons (≤15 nts) represent < 1% from the total number located in the coding genes in all genomes. Likewise, annotations of coding genes exhibiting intron sizes with an *excess* or *deficit modulo* 3 (*i.e.*, coding regions probably mistaken as introns and *vice versa*, respectively) are also unlikely to occur in our dataset (see Supplementary Figure S1 and Table S3). We acknowledge that particular phylogenetic patterns might be challenged by the completeness of genome assemblies in those lineages with very few sequenced genomes [66] or that will undergo substantial genome size corrections [128]. However, we also demonstrated to a considerable extent that possible biases in the genome assemblies and annotations analyzed here do not significantly impact our correlations and overall findings, after testing the genome completeness of 461 projects over 200 randomly reduced datasets. In disagreement with [66], thus, we show that the genomic differences obtained from most genome projects are robust enough to evaluate biological and evolutionary large-scale patterns of genome features across eukaryotes.

### Intron-richness is a suitable pre-condition to evolve complex multicellularity

Despite the many origins of multicellularity on Earth, complex multicellularity (CM) evolved only a few times in Eukarya [129–131]. We developed here a conceptual framework to define CM beyond the number of unique cell types (UCTs) (see Appendix 1). Our definition follows West and colleagues [132, 133] in acknowledging that contingent irreversibility from clonal-unitary development is key to evolve obligately multicellularity, but differs in arguing that a reinforced irreversibility of developmental commitment from multiple cell types is the main determinant in clonal multicellularity for a major transition to occur in individuality. By formally differentiating simple and CM, we thus hypothesize that CM is the outcome of *major evolutionary transitions* [133, 134] involving the presence of innovatory changes in genome structure and expression due to the differential evolution of particular ncDNA classes along the eukaryotic lineages [131]. We found here that complex multicellular organisms (CMOs) are characterized by high intron-richness; even those CMOs that have undergone strong selection to reduce several classes of ncDNA and genome size, such as carnivorous plants [135] and birds [76, 88, 89]. We show that the association between CMOs and intron-richness does not depends on changes of genome size, which in is agreement with the study of Niklas [136] indicating that increases in the number of UCTs fail to keep pace with increases in genome size. Our findings also suggest that CM origins were most likely preceded by high intron-richness, since the latter is also found on the closest unicellular and simple multicellular relatives of CM lineages. This is consistent with episodes of rapid and extensive intron gain found on the basal lineages of opisthokonts, holozoans and plants [1, 13, 137]. Furthermore, the diversity of intron-richness observed here among CMOs is not random nor homogeneous. Instead, it appears to be constrained by different factors that demand further research, including shared phylogenetic history, widely divergent selective regimes, lifestyles and generation times.

It is becoming clearer that intron-richness has important phenotypic consequences on eukaryotes, but which of these consequences can be considered indispensable to promote CM convergently? As summarized earlier, the functions –either causal roles or selected-effects– of introns promoting the emergence and evolution of CM can be very diverse. Most of these functions are, however, neither exclusive of CMOs nor universal across eukaryotes, but rather the outcome of exaptations originated on independent occasions [138]. This is partially because, as shown in this study, introns possess different characteristics throughout the major supergroups. Also, the rates of intron conservation and the molecular mechanisms responsible for intron processing vary considerably across eukaryotes [2]. Most importantly, introns appear to affect virtually every step of mRNA maturation, as described previously and reviewed in [138]. Yet, the role of exon skipping (ES) has been highlighted as the main promoter of multicellular complexity by expansion of proteome diversity [2, 59] through selection of, for instance, new transcription factor families, cell adhesion and signal transduction proteins [98, 139, 140]. However, most ES events and isoforms (mainly in low abundance) are found to be mainly the outcome of stochastic splicing errors [141, 142]. Also, transcriptome analyses from diverse tissues and cell lines reveal that most genes express one and the same dominant transcript in multiple tissues in human [143], mouse [144] and fly [145] (but see [146, 147]). It remains thus to be fully understood to what extent ES events are actually contributing to the suggested protein diversity of CMOs [59, 131].

The phenotypic diversity of CMOs largely relies on the expression of ancestral and species-specific genes coordinated in a particular spatiotemporal manner. We argue here that intron-richness has facilitated this process to a great extent by tuning the transcriptomes of an organism through intron-mediated mechanisms (IMMs) that alter the timing or kinetics of transcript expression. For instance, *intron retention* coupled to components of the RNA surveillance machinery can modulate gene expression post-transcriptionally by slowing splicing kinetics of those intron-containing transcripts stored at the nucleus in response to a variety of cellular signals [138, 148, 149]. Diverse modes of transcript regulation by intron retention have been found during cell type differentiation, cellular stress, circadian rhythm and early embryogenesis in plants [147, 150–152] and mammals [58, 153–156]. Alternatively, *intron delay* can coordinate the expression of genes that are sensitive to changes in their transcript length during particular stages of the metazoan development [33, 34]. The transcriptional delay caused by the presence of introns in the *Hes7* locus, for instance, controls the proper oscillatory expression of the genes involved in body segmentation during early vertebrate embryogenesis [36, 157–159]. Notably, the intron lengths of *Hes7* and of other genes involved in developmental patterning across mammals are highly conserved and even coevolving among coexpressed genes [160, 161]. These findings are consistent with studies showing that some species with long-complex life cycles and slowly regulated cell-cycle genes appear to be enriched in intron-rich gene-structures and patterning processes [33, 34, 148]. On the other hand, short life cycles and rapidly regulated cell-cycle genes (at different stages of development) tend to constrain gene-structures toward short and intron-poor genes for efficient expression [7, 26, 34, 35, 37, 69, 162, 163].

Rather than a selective (ultimate) effect, a major influence of IMMs to differentiate functional from nonsense transcripts in CMOs is expected if we consider that: (i) intronic RNAs constitute a considerable fraction of the transcriptomes in the CMOs analyzed [28–30]; (ii) canonical and non-canonical splicing errors (from which these IMMs emerge) appear to be more frequent when intron-richness increases [49, 58, 61, 141, 142]; and (iii) the spatiotemporal patterns of transcript expression derived from IMMs do have significant ecological and evolutionary consequences for cell cycle control and body plan formation [33, 34, 148, 149]. Nevertheless, some of the phenotypic consequences of intron-richness are also expected in certain life histories that did not evolve CM due to different evolutionary conditions. For instance, IMMs appear to influence the development of complex life cycles in intron-rich unicellular and simple multicellular species (as defined in Appendix 1) such as Apicomplexan parasites, which often involve multiple hosts and/or differentiation stages [96, 164]. Similar findings are expected to be found in the upcoming complete genomes of other CMOs within the red algae and *coenocytes* such as *C. taxifolia* [165]. Ultimately, major evolutionary transitions require extreme conditions for certain factors to become consistently important [133]. We argue that high intron-richness (through ES and IMMs processes) has laid the foundations for the emergence of novel mechanisms of transcriptome timing in Eukarya, which under exceptional conditions might have convergently co-opted for fate specification and commitment of cell types. It remains to be known whether common life-history traits [166, 167], molecular mechanisms [168, 169] and evolutionary forces have parallelly shaped the evolution of both intron-richness and tissue-based body plans in Eukarya.

## METHODS AND MATERIALS

### Genome-based data collections

We compiled the complete-sequenced nuclear genomes, protein-coding genes and gene annotation files for a total of 461 eukaryotic organisms from publicly available databases: 131 fungi, 78 species from Archaeplastida, 186 from Metazoa, 20 from Alveolata, 17 from Stramenopiles, 7 from Excavata, 7 from Amoebozoa, 4 from Choanozoa, 3 from Rhizaria, *Fonticula alba* (Fonticulidae), *Guillardia theta* (Cryptophyta), *Emiliania huxleyi* (Haptophyta), *Thecamonas trahens* (Apusozoa). A manual filter was applied to avoid redundant species (*i.e.*, same genus with similar genome sizes), sequenced genomes with <70% of the estimated genome size, and gene annotations without support from transcript data. To account for significant under and overestimations of genome contents, we also corrected our calculations with the “estimated” genome sizes based on experimental approaches collected from databases and literature. References and details of these datasets are provided in Supplementary Table S1.

### Estimating intron features with GenomeContent.pl

As depicted in Appendix 2-Figure 1, `GenomeContent` was written in Perl to calculate global statistics and sequence-based estimators of genome features through six major steps: (1) identifying coordinates from protein-coding gene (CDS); (2) checking the quality of intron annotations; (3) calculating statistic descriptors for genome-wide features from “reference gene sets” (derived from 1 and 2), such as size, density and number; (4) estimating genome-feature contents from the overlapping projection of “reference gene sets” onto the genome sequence; (5) calculating statistic descriptors for genome-feature contents; (6) plotting of figures and retrieving of sequences (fasta format) and statistics (text format). According to the definitions described in Appendix 2, we measured 30 genome features with `GenomeContent` across 461 eukaryotes, including 10 associated to the intron-richness of a genome: intron size (average and “weighted”, see equation 1 in Appendix 2), total number, absolute density, genomic content, number and fraction of genes containing introns. As observed in Appendix 2-Table 1, “(absolute) intron density” was measured as the average number of introns per intron-bearing gene because other estimations based on the average number of introns either per sequence region or from the total number of protein-coding genes are vulnerable to both genome size and considerable fluctuations of gene models, respectively.

### Determining the repeat content of genome features and statistical tests

We focused on identifying *de novo* repeats along the genome, rather than on classifying them in specific families. The `Repeatscout` algorithm v1.0.5 [170] was employed to compare a genome sequence against itself and in the two reading directions in order to identify *de novo* repetitive sequences with the minimal *k*-mer length of 15 nts. To gain major repeat coverage, the *de novo* `Repeatscout` libraries were merged with the Repbase libraries version 20.03 (http://www.girinst.org/repbase/). Then, these merged repeat libraries were used to map the coordinates of the repeats (interspersed and short repeats, and low complexity sequences) across the complete genomes with the `RepeatMasker` program v4.0.5 (www.repeatmasker.org). The placeholders were also taken into account as an independent category of “potential undefined repeats” within each genome feature (see Figure 2). However, the proportion of such regions in a genome is not collected in an individual genome feature for statistical analysis, given that they usually include the highly repetitive heterochromatic sequences that could not be unambiguously sequenced. Based on the genome coordinates obtained from this process, a nucleotide was classified as repetitive if it was covered by a repetitive sequence on either strand. Accordingly, the **repeat content** within introns, exons and the genome was determined as the total number (or percentage) of nucleotides in the genome feature that were classified as “repeat”; while the **unique content** of a particular genome feature is thus estimated from the non-repetitive nucleotides.

With a custom Perl script, we then calculated the *Jaccard index*, as *J*(*R, GF*) = |*R* ∩ *GF*|/|*R* ∪ *GF*|, to estimate the nucleotide overlap between the repeat coordinates (R) and the genome features (GFs: exons and introns) by using the “feature content coordinates” of every genome as observed in the Appendix 2-Figure 1. We tested the significant degree of observed overlap between repeats and GFs for each genome with the GenometriCorr package [171] from the R program v3.1.2 (www.r-project.org). For each genome, 1,000 permutations were allowed to shuffle the repeat coordinates along the genome sequence. Exon and intron positions were preserved as reference, while the repeat coordinates and the random sets were provided as the query. This setting provides the correct assessment of correlations (*p-value*) for the relative distance under the Kolmogorov-Smirnov criteria and for the permutation test on the *Jaccard index* [171, 172], which indicate whether the overlap is less (TRUE) or more (FALSE) significant than expected by chance. The statistical reports are available as Supplementary Data.

### Construction of phylogenetic trees

We constructed four different tree topologies, each of which has been extensively used in literature, for the 461 eukaryotes analyzed in this study, in order to: (a) evaluate phylogenetic signal, (b) perform phylogenetic controlled analyses, and (c) test the robustness of our comparative analyses against some of the phylogenetic uncertainty surrounding the tree of eukaryotes, owing to phylogeny construction errors and the absence of the true but unknown species tree.

To test for sensitivity to topological and branch length errors, we constructed a phylogeny determined by protein domain content as described in [173] (Figure 1d). Accordingly, *hmmscan* from `HMMER` v3.1b1 [174] was used to search for the protein domain models of the Pfam-A families from the PFAM database v30.0 [175] against the 461 proteomes. We used the gathering threshold (`–cut-ga`) for filtering out false positives. Custom `Perl` scripts were used to obtain a presence/absence matrix for all pfam domains detected and to calculate a pairwise distance matrix for all analyzed species. A weighting factor was included to correct the distance between two genomes (owing to the great differences in genome size, gene content, and lifestyles), and according to the following relationship: *D* = *A*^′^/(*A*^′^ + *AB*), where *A*^′^ is the number of unique pfam domains in one of two genomes compared: *A* and *B*, and *AB* is the number of pfam domains they share. Thus, “the two tendencies are acknowledged by setting the evolutionary distance equal to the ratio of the unique domains in the smaller genome (*A*^′^) to its total number of domains (*A*^′^ + *AB*)” [173]. The phylogeny construction was performed with the neighbor-joining method and bootstrapping with the program *neighbor* from `PHYLIP` v3.68 (http://evolution.genetics.washington.edu/phylip.html). We used *Trypanosoma brucei* as an outgroup, based on the supported basal phylogenetic position of Eozoa (Excavata and Euglenozoa) within Eukarya [176, 177].

Additionally, two NCBI taxonomy-based trees with no branch lengths were obtained with the species IDs collected from the “Taxonomy Browser” of NCBI (https://www.ncbi.nlm.nih.gov/Taxonomy/taxonomyhome.html/, last accessed July 14, 2016, see Supplementary Table S1) and with a combination of the phyloT (www.phylot.biobyte.de) and `iTOL` (www.itol.embl.de/itol.cgi) tools to allow or not the use of polytomies on each tree. A fourth tree topology (unrooted and with no branch lengths) was obtained by manually “correcting” the NCBI taxonomy-based tree (not polytomies allowed) with the `TreeGraph 2` v2.13.0-748 beta program [178] to fix the resolution at the genus and species level according to: (1) clade-specific phylogenies based on candidate orthologous sequences reported on literature, and (2) supertrees reconstruction for those few species that have not been incorporated into a sequence-based phylogeny yet. Polytomies were introduced in cases where phylogenetic uncertainty is not solved according to different studies. The 110 references employed to construct this consensus tree, some of which include the phylogenies reported along with the complete genome projects, are available on Supplementary Table S1. The literature consensus-based tree was selected as the “reference eukaryote tree” (Figure 1a) to present the results throughout the article. All tree topologies are available as Supplementary Data.

We quantitatively estimated the dissimilarity among the four trees with two measures reported to performed best among topology-only metrics [179]. The symmetric difference of Robinson and Foulds (RF) measures the number of different partitions (or clades not shared) between two trees [180], whereas the tree aligment metric (Align) of [181] scores the mismatches in the best alignment of the similar (and same) branches between two trees. Absolute RF distances were also divided by the total number of species in the tree to estimate the number of partitions per species. The Align and RF scores were calculated with the python scripts implemented in [179] and available at http://datadryad.org/resource/doi:10.5061/dryad.g9089.

### Phylogenetically corrected analyses

We first evaluated the strength of phylogenetic signal (*i.e.*, their statistical non-independence) exhibited by the 30 genome-based features analyzed in this study with the tree topologies described previously. Thus, we calculated the Pagel’s lambda (*λ*) transformation for all genome features analyzed with the *caper* `R` package (`pgls`) [182]. In comparison to other indices, Pagel’s *λ* is very robust to both incompletely resolved phylogenies and suboptimal branch-length information [83, 183]. We then calculated coefficients of determination (*r*^2^) to estimate the strength of the correlation to associate the variations observed between two traits (*X* and *Y*) with three linear regression models: Ordinary Least Squares (OLS) (*stats* `R` package: `lm`), Phylogenetically Independent Contrasts (PICs) (*ape* `R` package: `pic`) [184], and Phylogenetic Generalized Least Squares (PGLS) (*caper* `R` package: `pgls`). We also calculated log Bayes Factors (*LogBF*) to estimate the significance of evidence for the correlation between *X* and *Y* with the Markov chain Monte Carlo (MCMC) method and the PIC and PGLS models. log Bayes Factors were calculated as:

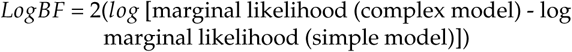

with 100 stones and 10,000 iterations per stone to estimate the marginal likelihood, as implemented in the `BayesTraits` program v3 (http://www.evolution.rdg.ac.uk/BayesTraitsV3/BayesTraitsV3.html).

We tested the robustness of the phylogenetically controlled regressions to discrepancies in tree topologies, phylogenetic diversity and estimations of genome features. First, OLS, PGLS and PIC regressions were performed with two sources for genome size: genome assemblies and experimental estimations. Also, the PGLS and PIC regressions were performed with the four tree topologies described previously. For the protein domain-based phylogeny, we also performed PGLS regressions with both equivalent branch lengths (*all* = 1) and lengths derived from the distance matrix. To test the influence of not fully resolved trees owing to the presence of hard polytomies, we generated three additional topologies for the NCBI-taxonomy tree (with polytomies allowed), two of them with randomly resolved polytomies using the procedure `multi2di=TRUE`, and one tree with a non-random procedure `multi2di=FALSE`, with the R package *ape* (library `picante`). Furthermore, we employed the “replicated co-distribution” approach [87] to test whether the association between *X* and *Y* is replicated across multiple independent clades. Thus, PGLS regressions were performed over 20 different sets compiled from the original dataset of 461 eukaryotes: 18 datasets correspond to divergent lineages, and two datasets (with 100 replicates each) were created through the random selection of 231 and 116 species, respectively. To further test the impact of phylogenetic diversity on the sensitivity of our phylogenetically controlled correlations, we performed PGLS regressions with four additional datasets (see Supplementary Table S12): two further randomly reduced datasets of 58 and 29 species, another dataset with 26 out of the 30 eukaryotes used in the study by Lynch and Conery [62], and the dataset of 30 eukaryotic genomes from the study of Wu and Hurst [68]. All genome-feature values were log10-transformed prior to analysis, except for the few genome-feature values == 0 that estimated the absence of repeats within introns in some extreme intron-poor genomes such as *Debaryomyces hansenii, Encephalitozoon cuniculi, Giardia intestinalis, Spironucleus salmonicida*. These particular values were discarded from the corresponding analyses.

We computed the Cronbach’s reliability coefficient alpha to measure the internal consistency (inter-relatedness) of the variables employed to test their relationship with multicellular complexity with the R package *psych* (`alpha`) [185]. Comparative and phylogenetic comparative Principal Component Analyses (PCA) were performed using the R packages *stats* (`princomp`) and *phytools* (`phyl.pca`), respectively. Branch transformations (*all* = 1) for the phylogenetically controlled PCAs and PGLS regressions were performed with *ape* (`compute.brlen`). Remaining statistical tests were calculated with custom R scripts. The plots were prepared with the R packages *ggplot, ggbiplot*, `phenotypicForest` v0.2 (http://chrisladroue.com/phorest/) and the software `Inkscape` (https://inkscape.org/en/). The results displayed throughout the paper are based on the estimated genome sizes and the “reference eukaryote tree” (Figure 1a). The results obtained with alternative tree topologies and assembled genome sizes are available in Supplementary Material.

## AVAILABILITY

The `GenomeContent` program and data are available upon request during peer-review, and will be openly available after publication.

## SUPPLEMENTARY DATA

Supplementary Data, Text, Figures and Tables are only available for peer-review at the moment, but will be openly available after publication.

We gratefully acknowledge the work and policies of the consortia and research groups for making publicly available the current wealth of genome datasets. We thank Jens Steuck and Ronald Kriemann for their assistance with the IZBI and MPI clusters required for computational work, A. Nabor Lozada-Chávez for his assistance with the R package, and Cristina Bojórquez-Espinosa for proofreading. We deeply appreciate the feedback received from previous anonymous reviewers.

### APPENDIX 1

#### Defining simple *versus* complex multicellularity

***Multicellularity*** refers to the phenotype characterized by the self-organization of cells that undergo a transition in individuality to perform cooperative consumption of energy, survival, and ultimately, reproduction. Multicellularity has arisen multiple times during the evolution of life on Earth [129], and it can even be induced in experimental settings [186, 187]. However, multicellularity is hypothesized to unfold into two different transitions [130, 131, 167]: *simple* or *complex*. Complex multicellularity (CM) is restricted to Eukarya and has evolved independently a few times: florideophyte red algae, laminarian brown algae, viridiplantae, eumetazoan animals, basidiomycota and ascomycota fungi. Typically, the number of unique cell types (UCTs) is used as the defining feature of multicellular complexity. However, accurate estimates of UCTs are only available for a small fraction of the species, and they also fail to appropriately capture the complexity of multicellular species [136]. Thus, we have created three “working definitions” embracing four criteria that distinguish few distinctive aspects of cellular development and life cycle to recognize species in our dataset, first as unicellular or multicellular (criterion 1), and then as simple multicellular (SMO) or complex multicellular (CMO) (criteria 2-4):

##### Criterion 1

*Whether there are one or several differentiated state cells at once across a life cycle.* Some single-celled organisms may have several differentiated state cells but at different times during the life cycle, such as the yeast *Saccharomyces cerevisiae* (three UCTs) [188], or may develop a *pseudo-hypha* with very few spores during a transient stage of a life cycle to reproduce [189]. Also some naturally living unicellular organisms, such as *S. cerevisiae* and *C. reinhardtii*, may develop filamentous growth under particular experimental settings that induce stress, and as long as the selective pressure is mantained [186, 187].

##### Criterion 2

*Whether the organization of the cells in a multicellular organism falls into one of the following states: a) differentiated cell types; b) undifferentiated cell types; or c) “syncytium/coenocyte” sensu amplio, i.e.*, multiple nuclei (either genetically identical or distinct) distributed within one common cytoplasm, which might or not be partially separated by cell membranes. Examples of the latter include: coenocytic *Dictyostelium discoideum* [190], siphonous algae and non-septate fungi [191]. Some coenocytic organisms, such as *Caulerpa taxifolia* [165], have morphological structures equivalent to a multicellular organ but not comprised in tissues or cells (*i.e., pseudo-organs*). While SMOs undergo a transition in individuality through any of the three cellular states, CMOs only undergo a major transition in individuality through differentiated cell types.

##### Criterion 3

*Whether the transition in individuality, as defined in criterion 2, is facultatively or obligately replicated across generations.* Facultatively multicellular species are able to complete their life cycle as unicells and only become multicellular under certain environmental conditions [132, 133]. For example, formation of fruiting bodies in some organisms, such as *Dictyostelium*, is observed in particular generations that undergo critical conditions to increase dispersal success [192, 193]. By contrast, obligately multicellular species can only complete their life cycle as multicellular organisms, owing mainly to the high genetic relatedness of cells originated through clonal-unitary development [129, 132, 133]. For instance, the development of tissue-based fruiting bodies (“basidia” and “ascocarp”) used for sexual reproduction in fungal CMOs is replicable on every generation. While SM is either facultatively or obligately replicated across generations, CM is only replicated as a whole. This criterion takes into account the temporal “unicellular transition” that all multicellular organisms, either simple or complex, undergo by means of reproductive processes through the life cycle [194].

##### Criterion 4

*Whether or not the transition in individuality, as defined in criterion 2, is produced by irreversible tissue-based body plans*. CMOs have tissue-based body plans that are developmentally irreversible, so that the within-group conflicts produced by “mutant-selfish” cell lineages (*defectors*) are negligible enough to avoid reversible differentiation of the whole organism [195–197]. Such irreversibility is consequence of active developmental commitment of multiple cell types that undergo fate specification and determination at particular stages of an organism life cycle. Cell type commitment is observed during the formation of: i) germ layers in metazoans [168] and eumetazoans such as cnidarians[198–200], ii) meristems in plants [201, 202] and in the CMOs within brown and red algae [203–205], and iii) in the primordium of fungal CMOs, although some cell types are still able to revert to vegetative growth *in vitro* [100, 206]. By contrast, SMOs do not develop true tissue-based body plans, owing in part to the lack of cell types with fixed identities and lineage commitment, so that dedifferentiation or transdifferentiation of cell types at any stage of development is common under the influence of certain factors. For instance, an absence of true tissue-based body plans, fate determination and stability of key cell types is observed in the sea sponge *A. queenslandica* (∼11 UCTs) [207, 208], and the sea placozoan *T. adhaerens* (∼5 UTCs) [209, 210] (but see [211, 212]). Likewise, the green algae *V. carteri* also lacks of a tissue-based body plan, since it only forms a colony of ∼2,000 cells with two UCTs [213].

According to these four criteria, we distinguish:

*Unicelullar:* is an organism exhibiting a single differentiated state cell at once across its life cycle. Single-celled organisms developing a transient *pseudo-hypha* or experimentally-driven filamentous growth are also included in this category.

*Simple multicellular*: is an organism exhibiting a facultatively or obligately transition in individuality through the organization of either several cells (with none or only few differentiated cell types) or a syncytium/coenocyte *sensu amplio* originated from one or more cell-line ancestors. Coenocytic and siphonous organisms structurated in *pseudo-organs* are also included in this category. Reversion to unicellularity may occur.

*Complex multicellular*: is an organism exhibiting an irreversible transition in individuality produced by tissue-based body plans, through the developmental commitment of multiple and different cell types originated from a common cell-line ancestor. Reversion to unicellularity or to a simple multicellularity lifestyle does not occur.

The lifestyle and body plan development of all species in our dataset were compiled from literature to evaluate the four criteria of our definitions, such information is available in Supplementary Table S2. We classified 77 species as unicellular, 96 species as SMOs, and 288 species as CMOs. This approach was useful to define the cellular state of some controversial model organisms. However, it still represents a challenge to distinguish between SM and CM in species within Fungi and Parazoa, whose multicellular body plans are generated from a few UCTs or are not well documented yet to evaluate the criteria described previously. Since this is a pioneer attempt to formally define CM and SM, we acknowledge that our definitions and the classification for the most controversial cases in this study are not free of future improvements and corrections.

**Appendix 2 Figure 1.**
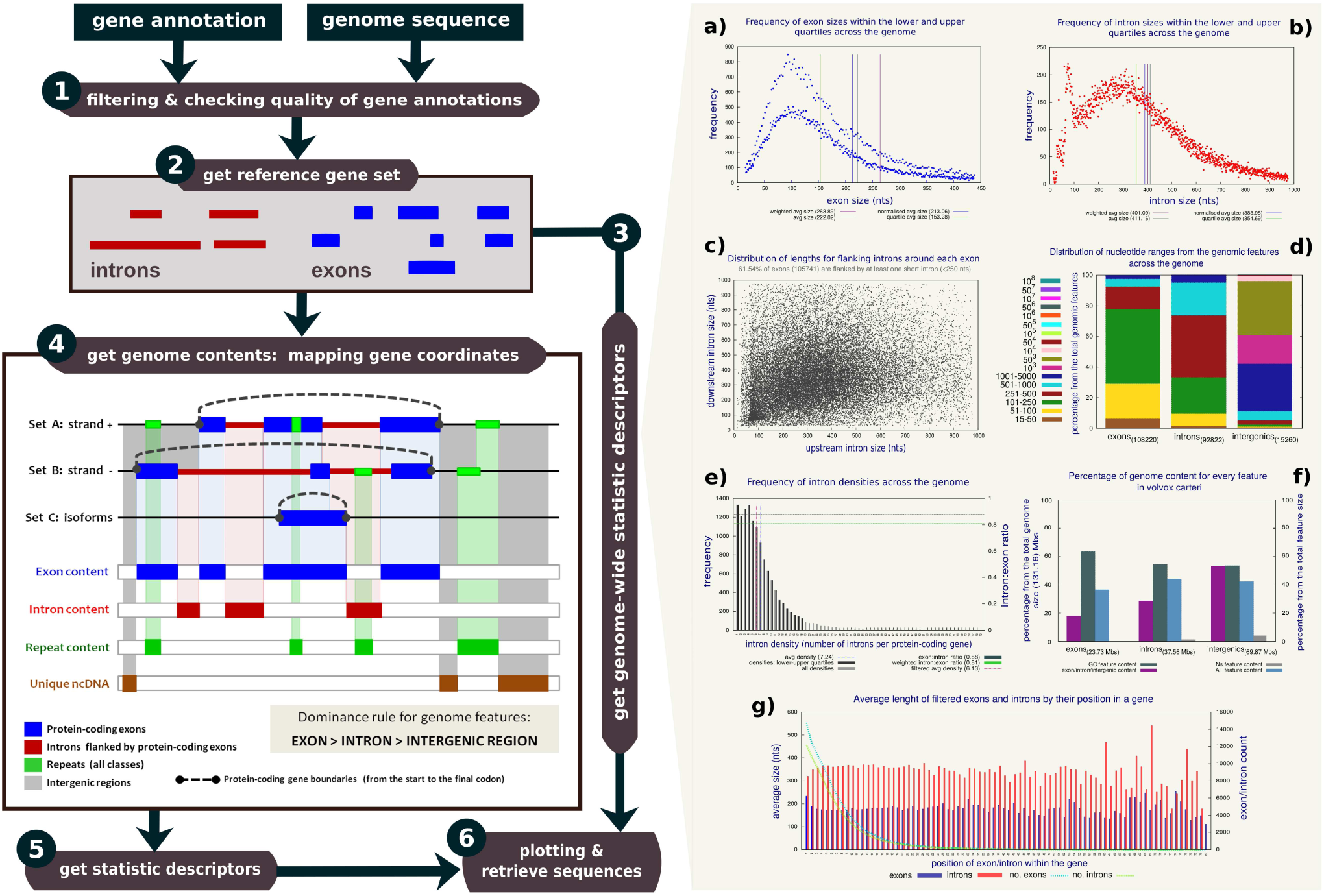
The GenomeContent program. A flowchart of the program is displayed on the left and described throughout Appendix 2. On the right, examples of exploratory figures showing some statistic descriptors for the genome of *V. carteri*, as obtained with the program.

### APPENDIX 2

#### A. Estimating intron features with GenomeContent.pl

`GenomeContent` was written in `Perl` to calculate global statistics and sequence-based estimators of genome features in six major steps, as shown in Appendix 2-Figure 1. First, the processing of gene annotations focuses on identifying coordinates from protein-coding gene (CDS), while the filtering process focuses on checking the quality of intron annotations. As described in next sections, the coordinates derived from both proceses are taken as the “reference gene sets” for introns, exons and intergenic regions to directly estimate several statistic descriptors, such as size, density and number. Then, the “reference gene sets” are projected onto the genome sequence in both strands, so that the nucleotide contents of each genome-feature are calculated according to the definitions described in a section below. Finally, all statistic descriptors obtained with the program are provided as text files, fasta formats and exploratory figures (see also Supplementary Figure S1). `GenomeContent` runs on an entire genome in few minutes or hours, depending on genome size and the number of annotated genes. `GenomeContent` is available upon request during peer-review, and will be openly available after publication.

#### B. Filtering of gene annotations

The filtering process of `GenomeContent` involves: a) identification of CDS, b) removal of small sizes, c) treatment of isoforms, and d) estimation of systematic errors in CDS. First, only genome coordinates from CDS were extracted, but their corresponding untranslated regions (UTRs) are not included because these are not fully annotated in most genome projects [214, 215]. Second, we excluded introns and exons smaller than 15 nucleotides (nts), which represent < 1% from the total number of introns and exons located within the coding genes of all genomes analyzed (as observed in Supplementary Figure S2 and Table S3). Third, alternative splice variants were kept in the data. To avoid redundant/overestimated data, however, exons with partial or full matching boundaries to exons of other transcripts were overlapped; the same rule was applied to introns. In both cases, their coordinates were joined or replaced accordingly; thus, every exon and intron is only counted once. We call this filtered set of protein-coding gene coordinates for every genome as the “reference gene set”.

**Appendix 2 Table 1.** Comparison of intron density estimations: absolute (aID) and normalized (nID).

| Species name | clade | aID | nID | # CDS | % CDS | # introns | genome size |
| --- | --- | --- | --- | --- | --- | --- | --- |
| *Tupaia chinensis* | Mammalia | 6.98 | 5.07 | 22,688 | 72.53 % | 114,940 | 3,200.0 Mbs |
| *Caenorhabditis elegans* | Nematoda | 5.28 | 5.12 | 20,520 | 96.81 % | 104,966 | 92.9 Mbs |
| *Gossypium raimondii* | Malvales | 6.00 | 5.20 | 77,267 | 86.86 % | 402,768 | 880.0 Mbs |
| *Xenopus laevis* | Amphibia | 7.41 | 5.25 | 43,025 | 70.81 % | 225,747 | 3,110.0 Mbs |
| *Guillardia theta* | Cryptophyta | 6.74 | 5.28 | 24,840 | 78.29 % | 131,044 | 87.2 Mbs |
| *Postia placenta* | Basidiomycota | 5.94 | 5.54 | 17,173 | 93.22 % | 95,089 | 90.9 Mbs |
| *Bombus terrestris* | Arthropoda | 6.25 | 5.99 | 8,334 | 95.80 % | 49,907 | 274.0 Mbs |

Most of the collected gene annotations are based on transcript evidence. Nevertheless, we implemented the approach proposed by Roy and Penny [216] in `GenomeContent` to further estimate systematic errors in CDS annotations by identifying the excess/deficit of the intron-length distributions modulo 3. Since introns are not expected to respect the coding frame, intron lengths 3*n*, 3*n* + 1, and 3*n* + 2 should appear in similar fractions *p*3*n* ≈ *p*_3*n*+1_ ≈ *p*3*n*+2. As stated in [216], large values of “3n excess”, *E*3 = *p*3*n* - (*p*_3*n*+1_ + *p*3*n*+2)/2, suggest that a considerable fraction of internal exons may have been incorrectly predicted as introns or that there are several “intron retention” events. On the other hand, a deficit of 3*n* introns, i.e., *E*3 ≪ 0, suggests that a considerable fraction of 3*n* introns –lacking of stop codons– may have been mistaken for exons. Most gene annotations included in this study shown values of the 3n excess close to 0. Very few genomes (such as parasites and endosymbiots) were initially excluded from the present study because they exhibited high 3n excess (0.4 - 0.7) (see Supplementary Figure S2 and Table S3).

#### C. Estimation of average sizes and density of introns

Several statistical estimators for every genome feature were obtained with `GenomeContent`. **Genome size** is defined as the net length of nucleotides and placeholders of all sequences conforming the nuclear genome. The average *feature size* (*e.g., intron size* and *exon size*) of CDSs per genome was calculated in two ways. The **straight average size** (*A*_*f eature*_) is calculated as the total length of all feature sequences (exons or introns) in a genome (*L* _*f eature*_) divided by the total number of all feature sequences (exons or introns, respectively) in a genome (*N*_*f eature*_): *A*_*f eature*_ = *L* _*f eature*_/*N*_*f eature*_. The *straight average* depends on the number of data points from the whole sample (*i.e.*, gene models with introns), which equally contribute to the final average regardless of which gene they belong to.

If we now consider *a* _*f eature*_ to be the average length of the respective feature (exon or intron) within one single gene: *a* _*f eature*_ = *l* _*f eature*_/*n* _*f eature*_, then the **weighted average size**, *ā* _*f eature*_, is calculated as the mean of the *a* _*f eature*_ values of the respective feature (exons or introns) in a genome, according to:

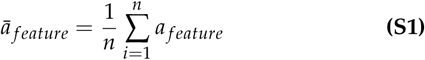

where *n* represents the total number of CDSs in a genome when calculating *āexon*, or the total number of intron-containing CDSs in a genome when calculating *ā*_*intron*_ [217]. The *weighted average* depends on the gene-structure of the genome, and thus it samples more broadly the data points that contribute, in the case of introns, to the well known skewed length distribution.

`GenomeContent` also estimates the abundance of introns within a genome with two different estimates to detect small changes of intron-richness and to buffer dramatic changes among updated genome releases and gene annotations. On the one hand, the abundance of introns across CDSs was estimated as the **fraction of intron-containing CDSs** from the total number of CDSs (% CDS), which might reflect complete intron loss from CDS structures at the genome level. On the other hand, the abundance of introns and exons within CDSs (“absolute density”) was estimated as the mean number per genome of exons in CDS (**exon density**), or introns per intron-containing CDSs (**intron density**), respectively. We employ the “absolute density” because, as observed on Appendix 2-Table 1, the average number of introns either per sequence region or from the total number of genes (“normalized density”) depends on both genome size and the number of CDSs, respectively.

For instance, the four species listed in Appendix 2-Table 1 exhibit around five introns per CDS when a “normalized” density is estimated from the total number of CDSs. However, the “absolute intron density” clearly shows that, for instance, *X. laevis, T. chinensis* and *G. theta* have indeed more introns per CDS on average than the other species, despite of having a smaller fraction of intron-containing CDSs (70.8%, 72.5% and 78.3%, respectively), and lower number of introns and CDS in some cases. Clearly, the bias observed in the “normalized intron density” is produced by larger numbers of total CDS in the genome.

##### D. Estimation of genome contents

`GenomeContent` also estimates the “feature content” of a given genome, *i.e.*, the proportion of nucleotides of the respective genome features (intron, exon, or intergenic) that contributes to genome size. Since most genome annotations only contain protein-coding regions rather than full transcript models, we count only coding exons and introns delimited by a pair of coding exons. As shown in Appendix 2-Figure 1, the program projects the sets of genomic intervals for all exons and introns from coding genes located in the plus strand (set *A*), the minus strand (set *B*), and of the isoforms (set *C*). Since a given nucleotide may be classified differently for different isoforms, we used the following domincance rule in order to obtain a unique classification at the genome level:

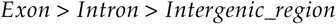

It reflects the idea that a genomic position is exonic whenever it appears in a coding exon of at least one transcript. Thus, the **exon content** of a genome is calculated as the total number of nucleotides in the genome sequence that are classified as coding exon with respect to at least one isoform. Analogously, a position is classified as ‘intronic’ if it appears inside the boundaries of annotated coding exons, but it does not overlap with any coding sequence. Thus, **intron content** was determined as the total number of nucleotides of a genome that were classified as intronic. The **CDS content** of a genome is calculated as the total number of nucleotides covered by the intronic and exonic positions within coding genes. Finally, the **non-coding DNA content** was computed analogously as the total number of nucleotides in a genome that are not covered by exonic and intronic positions from coding genes. Genome-feature contents are reported as: total size in Megabases (Mb), fraction (%) from the total genome size, and as genomic coordinates.

